# Dual localization of JA receptor, CaCOI2, explains JA perception dynamics in chickpea

**DOI:** 10.1101/2023.12.18.572114

**Authors:** Ajit Pal Singh, Ekampreet Singh, Amit Kumar Singh, Urooj Fatima, Muthappa Senthil-Kumar, Jitender Giri

## Abstract

Jasmonates (JAs) form a group of oxylipin-derived phytohormones involved in multiple biotic and abiotic stress responses and regulate plant development. JAs are perceived by the receptor proteins, COI (coronatine insensitive). These JA receptors encoding F-box protein, form SCF^COI^ ubiquitin ligase complex (Skp, Cullin, F-box) and activate JA signaling by facilitating degradation of the transcriptional repressor (JAZ; Jasmonate associated ZIM domain containing) proteins through 26S proteasomal pathway. However, JA signaling is poorly understood in chickpea, a vital legume plant. Here, we first identified two putative chickpea JA receptor named CaCOI1, CaCOI2 and demonstrated CaCOI2 as a functional JA receptor. Further subcellular localization studies revealed that CaCOI2 is localized to extra-nuclear region and move to the nucleus on JA perception to activate JA signaling. Using domain swapping (between CaCOI1 and CaCOI2) experiments, we have demonstrated that the LRR region of JA receptors, which interact with bioactive JA i.e., JA-Ile, also plays a critical role in regulating subcellular localization of CaCOI proteins. We report that an acidic amino acid in F-box region of COI proteins (AtCOI1^T29^) stabilizes COI-InsP8 (1,5-bisdiphosphoinositol-1D-myo-inositol (2,3,4,6) tetrakisphosphate) interaction. This interaction is found to be critical for COI-JAZ interaction and therefore, their functionality. Our study thus identified a functional JA receptor in chickpea and revealed novel aspects of JA signaling and perception, which might also have implications in other plants.

## Introduction

Jasmonates (JAs), a collective term representing jasmonic acid and its derivatives, have been established as a key plant growth and response regulator under various biotic and abiotic stresses (Wasternack and Hause, 2013; Wasternack, 2014; Singh et al., 2015; Huang et al., 2017; Zhai et al., 2017; Singh et al., 2020, Singh et al., 2021). Mutants defective in JA biosynthesis like *dde1* (Sanders et al., 2000); *opr3* (Stintzi and Brouse, 2000); *dad1* (Ishiguro et al., 2001; *aos* (Park et al., 2002); *acx1* (Schilmiller et al., 2007); *opr3-3* (Chini et al, 2018), and signalling *coi1* (Feys et al., 1994; Ellis and Turner, 2002) show defective reproductive development including shorter anther filament and impaired pollen development resulting into male sterility. Among the many conjugates of JA, JA-Ile (JA-Isoleucine) is the bioactive form that elicits JA signaling (Fonseca et al., 2009). JA is perceived by COI1 (CORONATINE INSENSITIVE 1, in Arabidopsis), named after a phytotoxin produced by *Pseudomonas syringae* (Feys et al., 1994; Yan et al., 2009). COI1 encodes the F-box protein component of SCF^COI1^ (Skp, Cullin, F-box) E3 ubiquitin ligase complex (Xu et al., 2002). The receptor interacts with JAZ (JASMONATE ZIM DOMAIN family) proteins in the presence of JA-Ile or COR (coronatine) and ubiquitinates these transcriptional repressors (Xu et al., 2002; Chini et al., 2007; Thines et al., 2007). This poly-ubiquitination of JAZ proteins marks their degradation through 26S proteasomal pathway, releasing bHLH (basic helix-loop-helix) transcription factors like MYC2 to activate the transcription of downstream JA-responsive genes. JAZ proteins have N’-terminal TIFYXG motif facilitating homo and hetero-dimerization of JAZ proteins among themselves and a C’-terminal jas motif involved in the interaction with COI1 and MYC2 like bHLH TFs (Wasternack and Hause, 2013). Apart from the TIFYXG and Jas motifs, a short loop region in JAZ proteins is also important for JA-Ile or COR binding and facilitating the ternary complex formation, which includes JAZ, JA-Ile and COI1 (Melotto et al., 2008; Sheard et al., 2010).

COI proteins are composed of N’-terminal F-box domain and a C-terminal leucine-rich repeats (LRRs). Also, 33 amino acid residues in COI proteins are known to participate in the interaction involving JAZ and COI1 protein in the presence of JA-Ile and inositol pentakisphosphate (InsP5) or inositol tetrakisphosphate (InsP4) (Sheard et al., 2010; Lee et al., 2013; Qi et al., 2022). Later, it was determined that higher InsPs like InsP6, InsP7 and InsP8 also interact with COI proteins. Further, *atvih2* mutant, deficient in InsP8, though had higher JA/JA-Ile content upon wounding but underwent lower JA signaling upon biotic infection than Col-0 (Laha et al., 2015). These results suggested an important role of InsPs in JA signaling or JA perception. Previously, in a separate study, seven more amino acid residues from LRR region were found necessary for coronatine/JA-Ile perception, suggesting that the LRRs of COI1 are highly critical for JA perception (Yan et al., 2009). Now, despite having resolved the structure of COI1, the mechanistic aspect of JA perception can only be hypothesized. Among the two prevalent theories for JA perception, one supports that the receptor COI1, binds to the ligand (JA-Ile or COR) first and then the target JAZ proteins (Yan et al., 2009; Yan et al., 2018), while the second theory supports the involvement of co-receptor complex (COI1-JAZ) in JA perception (Sheard et al., 2010). The latter leaves us with a question of whether the coronatine/JA-Ile is perceived by the co-receptor complex consisting of COI1 and JAZ. If yes, then why is the ligand (JA-Ile or COR) required to make these two proteins interact? The requirement of the ligand for the interaction itself suggests that the ligand interacts first with the receptor, then substrates are recognized, or the three of these interact simultaneously. Further, pre-incubated COI1 (with either JA-Ile or COR) was found to interact with JAZ1, while JAZ1 alone was neither able to bind with JA-Ile/COR nor with COI1 (Yan et al., 2018). Also, the detection of COR associated only with COI1 but not with JAZ1 supported JA perception by a single molecule i.e., COI1 (Yan et al., 2018).

In continuation of our previous work (Singh et al., 2015), we identified two potential JA receptors, CaCOI1 and CaCOI2 in chickpea. Interestingly, only one (CaCOI2) of the two identified JA receptors was found functional despite having all the previously known conserved amino acids in both proteins, revealing that additional amino acid residues still need to be identified. We observed that JA receptors follow a peculiar dual subcellular localization dictating their functionality. The two JA receptors from chickpea are localized along the plasma membrane while their targets are localized to the nucleus. We have demonstrated that JA receptors move to the nucleus on JA treatment, validating the single receptor model for JA perception from Yan et al. (2018). Utilising domain swapping strategies combined with subcellular localization and complementation assays, we validated that the C’-terminal LRR region is responsible for the nuclear localisation of JA receptors. Further, in-silico docking analysis was utilised to prove that an additional amino acid of acidic nature is essentially required for InsP8 interaction which further facilitate interaction of COI proteins with target JAZ proteins in the nucleus.

## Results

### The chickpea genome encodes two AtCOI1 homologs

Our previous report identified one CaCOI1 as an AtCOI1 homologue in chickpea (Singh et al., 2015). Since, the depth and coverage of chickpea genome sequence have significantly increased in the recent past, hence, we re-analysed the presence of AtCOI1 homologues in chickpea genome of both Desi (http://nipgr.res.in/CGAP/home.php) and Kabuli (http://www.icrisat.org/gt-bt/ICGGC/homepage.htm) cultivars. Here, we could identify two AtCOI1 homologues i.e., CaCOI1 and CaCOI2. Protein sequence analysis confirmed the presence of F-box domain and LRRs in the identified chickpea COI proteins using SMART (http://smart.embl-heidelberg.de/), Interpro (http://www.ebi.ac.uk/interpro/) and NCBI CDD (https://www.ncbi.nlm.nih.gov/Structure/cdd/wrpsb.cgi) search. The protein sequence alignment of CaCOI1 and CaCOI2 with AtCOI1 suggested that the F-box and LRR regions were highly conserved in newly reported COI proteins (Sheard et al., 2010; Qi et al. 2022). We also observed that the amino acid residues playing a critical role in the interaction between COI and COR or COI and JAZ were conserved in all three proteins analysed (Figure S1, Sheard et al., 2010; Lee et al., 2013; Qi et al., 2022). These observations suggested that both CaCOI proteins may act as functional JA receptors.

Phylogenetic analysis revealed that being dicot, CaCOI proteins were evolutionary more closure to Arabidopsis COI1 protein than monocot (rice and maize) COI proteins (Figure 1A). Further, gene structure analysis with GSDS (Gene Structure Display Server, http://gsds.cbi.pku.edu.cn/) revealed that despite the difference in the gDNA (genomic DNA) length of these genes, the coding region (CDS) and protein sequences were of almost equal lengths. The exon numbers and lengths were similar for Arabidopsis and chickpea COI proteins. However, we observed long intronic regions in both chickpea COI genes, while Arabidopsis COI gene had relatively shorter intronic regions (Figure S2).

**Figure 1.**
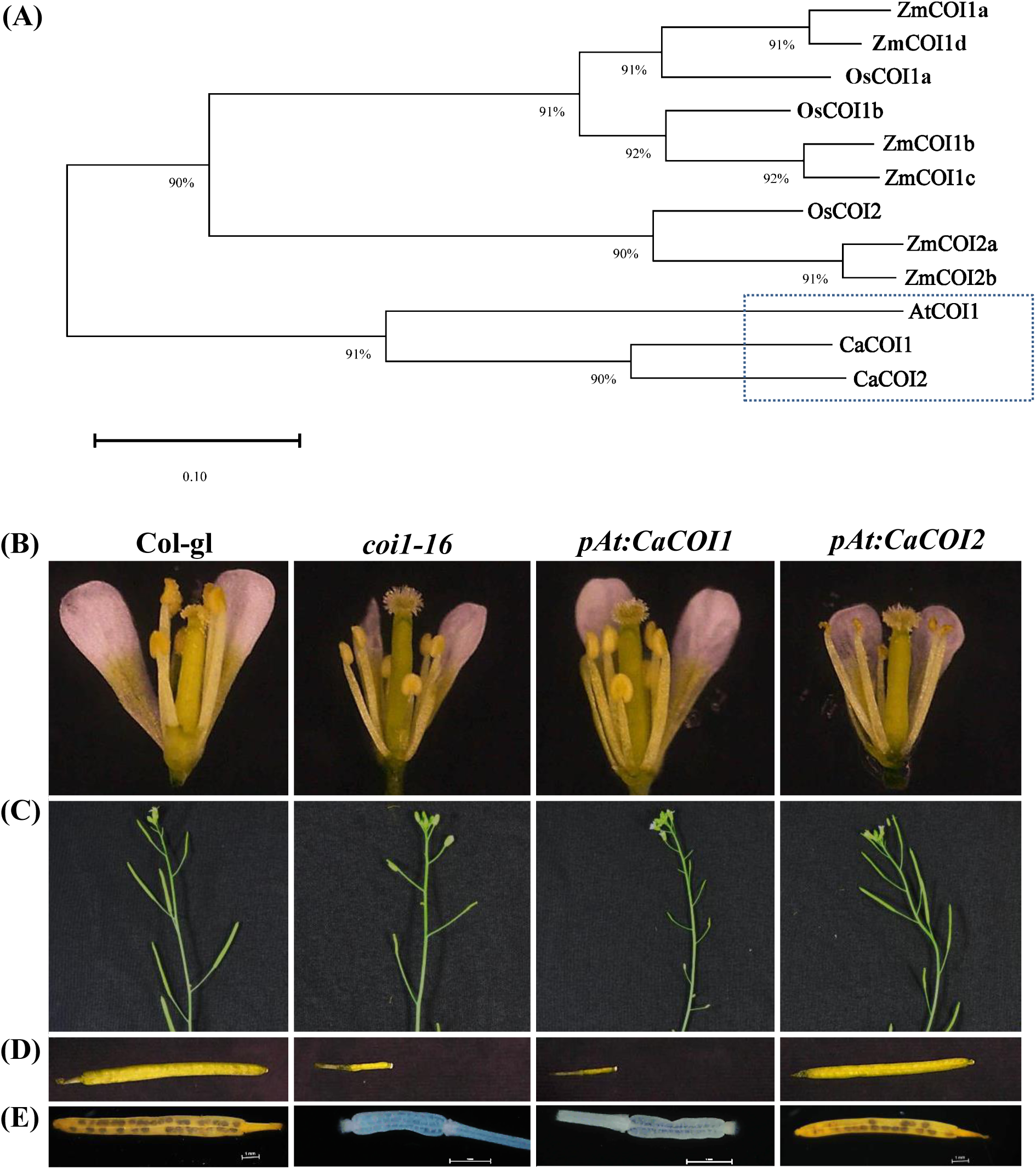
CaCOI2 complements *coi1-16* mutant. **A)** Phylogenetic tress with amino acid sequences of COI proteins from maize, rice, Arabidopsis and chickpea. CaJAZ6 protein sequence was taken as outlier. Peptide sequences were aligned using ClustalX and tree was generated using NJ method with maximum bootstrap value =1000. **(B-E)** Phenotypic complementation of *coi1-16* mutant with *pAtCOI1:CaCOI1 (pAt:CaCOI1)* or *pAtCOI1:CaCOI2 (pAt:CaCOI2)* using floral-dip method. **B)** Images showing floral morphology and anther dehiscence, **C)** silique formation, **D)** enlarged view of silique and **E)** seed development in siliques produced in Col-gl, *coi1-16*, *pAt:CaCOI1* and *pAt:CaCOI2* complemented *coi1-16* mutant. KI-I_2_ staining was performed to visualize seed development in siliques.

The whole genome sequence data for Kabuli genotype was available for CDC Frontier, a Canadian Kabuli chickpea genotype (Varshney et al., 2013) while for this study, we used an Indian cultivar, JGK-3. To identify any genotypic variation due to genetic diversification among the two genotypes, we analysed the coding sequence (CDS) of *CaCOI* genes from the JGK-3 cultivar. We observed 4 SNPs in CDS region of CaCOI1 (G at 376 position is converted to A (G376A), A570G, T638C and T1239C) and one SNP in the CDS region of CaCOI2 (C96G). In both genes, none of the SNP resulted in the premature termination of translation (Figure S3A). In CaCOI1, two SNPs were nonsense, while the other two resulted in A126T and L213P replacements. These two changes in a protein sequence are not expected to influence the interaction of COI and JAZs because these amino acids are not among the ones involved in such interaction (Sheard et al., 2010; Lee et al., 2013). In CaCOI2, the SNP changes remain nonsense and do not influence the protein sequence (Figure S3B).

### *CaCOI2* but not *CaCOI1* complements Arabidopsis *coi1-16* mutant

Arabidopsis *coi1-16* mutant fails to produce siliques at 22 °C but produces seed at 16 °C. We utilized this phenotype for functional characterization of identified CaCOI proteins. We transformed *coi1-16* mutant with *CaCOI1* and *CaCOI2 CDS* driven by *AtCOI1* promoter (Figure S4A-B). Seed formation was observed as the marker of complementation in *pAtCOI1:CaCOI1* (hereafter indicated as *pAt:CaCOI1*) and *pAtCOI1:CaCOI2* (*pAt:CaCOI2*) complemented lines of *coi1-16* at 22 °C. Surprisingly, none of the *pAt:CaCOI1* transformants were able to form silique at 22°C, while six lines of *pAt:CaCOI2* formed siliques similar to Col-gl (Figure 1B; Table S2). When open flowers of Col-gl, *coi1-16* and complemented lines were observed for visual markers of sterility, we found shorter stamen filaments in *coi1-16* mutant, which failed to reach stigma to fertilize it. Similar shorter filaments were also observed in *pAt:CaCOI1* transformant lines, while filament length was comparable in Col-gl and *pAt:CaCOI2* complemented lines (Figure 1B). Additionally, we observed highly intact anther in *coi-16* mutant and *pAt:CaCOI1* transformant lines, while anther dehiscence was visible in Col-gl and *pAt:CaCOI2* complemented lines. These two factors might have contributed to the male sterility of *coi1-16* mutant, but in the case of *pAt:CaCOI2* complemented lines, the stigma can now be fertilized and therefore, they can now develop siliques and seeds as produced by Col-gl (Figure 1C-E).

### CaCOI2 restores JA sensitivity in *coi1-16* mutants

Exogenous MeJA treatment is known to inhibit Arabidopsis root growth while *coi1-16* is relatively insensitive to MeJA treatment (Feys et al., 1994; Ellis and Turner, 2002). Therefore, we analysed the growth behaviour of Arabidopsis Col-gl, *coi1-16* and *pAt:CaCOI2* complemented lines on MeJA treatment. For this, we transferred 7-days-old ½ MS-Agar grown seedlings to 100 μM MeJA containing ½ MS plates and analysed the root growth inhibition after 7 days of the treatment. The primary root length inhibition assay suggested nearly complete restoration of MeJA sensitivity in *pAt:CaCOI2* plants compared to *coi1-16* mutant (Figure 2A-B; Figure S5A-B). Also, there was 12.5% and 27% inhibition in lateral root length of Col-gl and *pAt:CaCOI2* complemented lines, respectively, on MeJA treatment but, only 3% inhibition was observed in *coi1-16* mutant during the same conditions (Figure S5C). We also observed highly restricted root hair growth (number and length) in *coi1-16* mutant while Col-gl showed well-developed root hairs under normal growth conditions. Consistent with other phenotypic observations, all the complemented lines showed root hair development similar to Col-gl (Figure 2C). Further, anthocyanin accumulation was also found to be associated with Col-gl and complemented lines but not with *coi1-16* on MeJA treatment (Figure 2D). To investigate the restoration of JA signalling at the transcription level, we subjected one-month-old Col-gl, *coi1-16* and *pAt:CaCOI2* complemented lines (L2 and L6) to 100μM MeJA foliar spray (DMSO as control) and analysed the expression pattern of known JA inducible genes. Expression profiling showed the induction of *OPR3*, *AOS* and *VSP1* genes in Col-gl as well as complemented lines, while there was no induction of these genes in *coi1-16* mutant, confirming the restoration of JA signaling in the *CaCOI2* complemented lines (Figure S5D). We further tested *coi1-16* complementation on nutrient deficiency response by subjecting Col-gl, *coi1-16* and the complemented lines to phosphate deficient media. In agreement to the previous reports, we also observed *coi1-16* mutant relatively tolerant while Col-gl and the complemented lines showed higher sensitivity to phosphate deficiency mediated growth inhibition and senescence (Figure S6).

**Figure 2.**
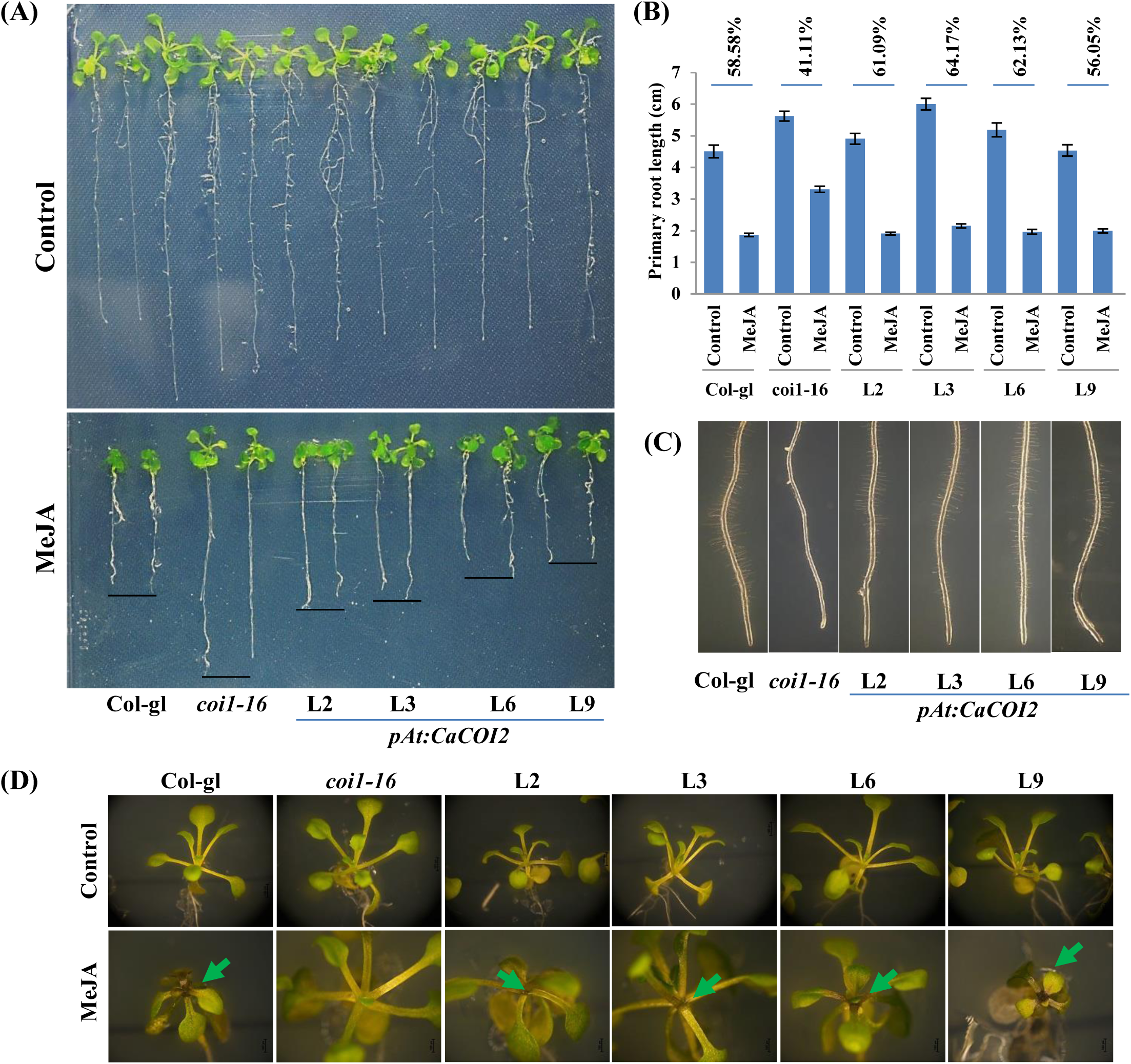
*CaCOI2* restores MeJA sensitivity to *coi1-16* mutant. **A)** Representative images showing root growth pattern in Col-gl, *coi1-16* mutant and *pAt:*CaCOI2 complemented lines under normal and 100µM MeJA treatment. **B)** Quantitative analysis showing root growth inhibition after 100µM MeJA. Each bar represents average of at least 30 biological replicates with SE among the replicates. **C)** Root hair development was analyzed using sterio-zoom microscope in 7 days old normally grown (1/2MS media) seedlings before transferring them to MeJA stress. **D)** Anthocyanin accumulation was observed under normal and MeJA treatment conditions using sterio-zoom microscope after seven day of treatment.

Lastly, AtCOI1 was named <u>Co</u>ronatine Insensitive because a mutation in this protein fails to recognise coronatine (a bacterial phytotoxin, structural mimic of JA-Ile) produced by *P. syringae* and therefore, its mutation imparts tolerance against *P. syringae* infection (Feys et al., 1994). To confirm CaCOI2’s role as a functional JA receptor, we examined host pathogen *P. syringae* infection and disease expression patterns in Col-gl, *coi1-16*, and complemented lines. As expected, the *coi1-16* mutant displayed reduced sensitivity to the infection, while the complemented lines exhibited sensitivity similar to Col-gl (Figure 3). Notably, bacterial pathogen multiplication in CaCOI2 lines closely resembled that of wild-type plants and was significantly higher than in the mutant. This consistency was evident through both conventional CFU quantification and the visualization of GFPuv-labeled pathogen in the apoplasts. Additionally, the disease expression phenotype and disease index indicated greater disease expression, characterized by chlorosis and cell death, in the CaCOI2 lines, akin to the wild-type, while mutants displayed minimal disease symptoms. Together, these results made us conclude that *CaCOI2* is the functional homolog of *AtCOI1* in chickpea.

**Figure 3.**
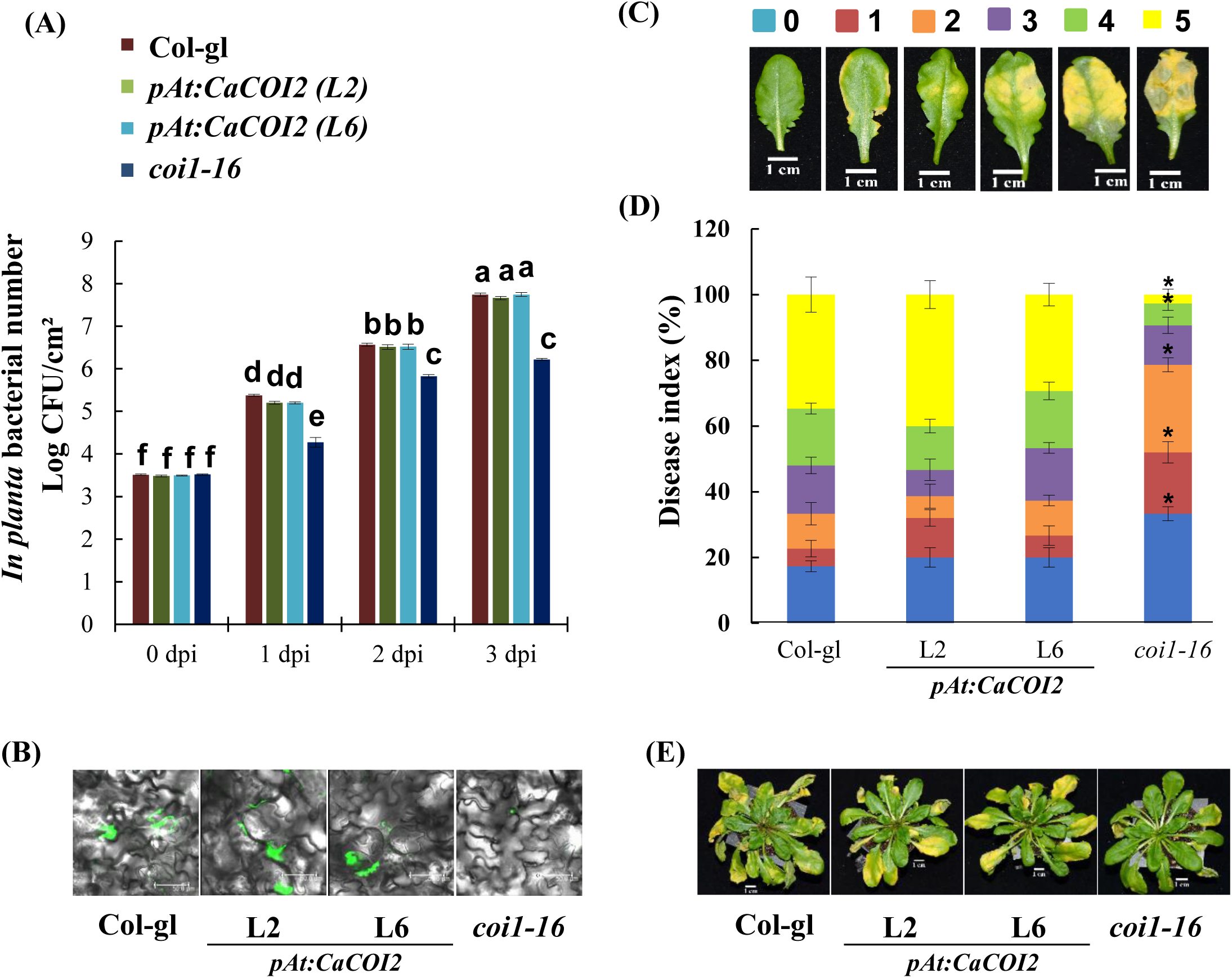
Complementation of *coi1-16* mutant with *CaCOI2* restores the susceptibility towards *Pst* DC3000. (A) Fully expanded leaves of Arabidopsis wild-type (Col-gl), complementation lines pAt:CaCOI2(L2) and pAt:CaCOI2(L6), and *coi1-16* mutant were inoculated with *Pst* DC3000 at a concentration of 5 X 10^5 CFU/mL. A two-way ANOVA test with Tukey correction (P < 0.05) was conducted, and significant differences are denoted by distinct letters. The bar represents the average of six biological replicates, with error bars indicating standard errors (SE) among the replicates. "Dpi" denotes days post inoculation. (B) *In planta* bacterial population of green fluorescent protein (GFP)-labeled *Pst* DC3000 was assessed at 2 dpi. Arabidopsis Col-gl, complementation lines, and mutant plants were inoculated with *Pst* DC3000 expressing GFPuv at 5 X 10^5 CFU/mL. At 2 dpi, leaf samples were examined using a confocal microscope (Leica TCS SP8). (C) Disease scores were assigned to monitor symptom progression. A score of 0 indicated leaves without symptoms, 1 for leaves with chlorosis at margins, 2 for leaves with chlorosis near the mid-rib region, 3 for leaves with two-thirds of the area affected by chlorosis, 4 for full leaves showing chlorosis, and 5 for leaves with both chlorosis and necrosis. (D) The disease index was determined based on disease scores for pathogen-infected plants at 3 dpi. Asterisks indicate a significant difference from the wild-type (Student’s t-test; *P < 0.01). Each bar represents the average of five replicates, with SE indicated among the replicates. (E) Phenotype of pathogen-inoculated plants observed at the 3 dpi stage.

### CaCOI2 but not CaCOI1 interacts with CaJAZ proteins

To regulate the JA signalling, COI proteins (in the presence of JA-Ile or coronatine) must interact and ubiquitinate JAZ proteins to facilitate their degradation via 26S proteasomal pathway. To investigate this attribute and consider upgradation of chickpea genome data, we identified potential JAZ proteins using the HMM profile and criteria used previously for the identification of JAZ proteins (Singh et al., 2015). This time, we identified 14 potential *CaJAZ* candidate genes, while previously, only 10 *Ca*JAZ genes were identified. All the *CaJAZ* genes were renamed according to their protein homology with Arabidopsis JAZ candidates (Figure S7; Table S3).

While analysing the genotyping variations, we did not observe any change in CDS of 7 *CaJAZ* genes, including *CaJAZ1b*, *-1c*, *-6*, *-9a*, *-9d*, *-9e* and *-12a,* while in 7 genes, either INDEL or SNPs or both were observed (Figure S8). However, none of these changes lead to a stop codon or loss of a functional domain. Therefore, genes in both genotypes encode potentially functional JAZ proteins. As JAZ proteins are known to form homo and hetero dimers, we analysed their interactions using one-to-one Y2H (yeast-two-hybrid) assays and found many of them forming homo and hetero dimers (Figure S9; Table S4). Now, considering them as functional JAZ proteins, we were interested to know whether CaCOI1 and CaCOI2 interact with JAZ proteins. One-to-one interaction using Y2H assay among CaCOI1/CaCOI2 and CaJAZs revealed no interaction between any of the CaCOI proteins and JAZ proteins in the absence of coronatine (Figure S10). However, CaCOI2 showed interaction with CaJAZ1a, 1c, 3b, 3c and CaJAZ12a in the presence of coronatine. Interestingly, CaCOI1 could not interact with any of the JAZ proteins, even in the presence of coronatine (Figure 4A). Secondly, the interaction of JAZ proteins with CaCOI2 also depends on the concentration of coronatine as we found a progressively increasing number of interactors when we increased coronatine concentrations (Figure 4A). Third, the entire CaJAZ9 clade failed to interact with CaCOI1 or CaCOI2. Amino acid sequence analysis of these proteins suggested that they have single amino acid residue commonly missing in the loop region (Figure 4B), which is known to be critical for interaction of JAZ proteins with JA receptors (Sheard et al., 2010). These observations further validated that CaCOI2 and CaJAZs are functional proteins while *CaCOI1* encodes for a non-functional JA receptor in chickpea.

**Figure 4.**
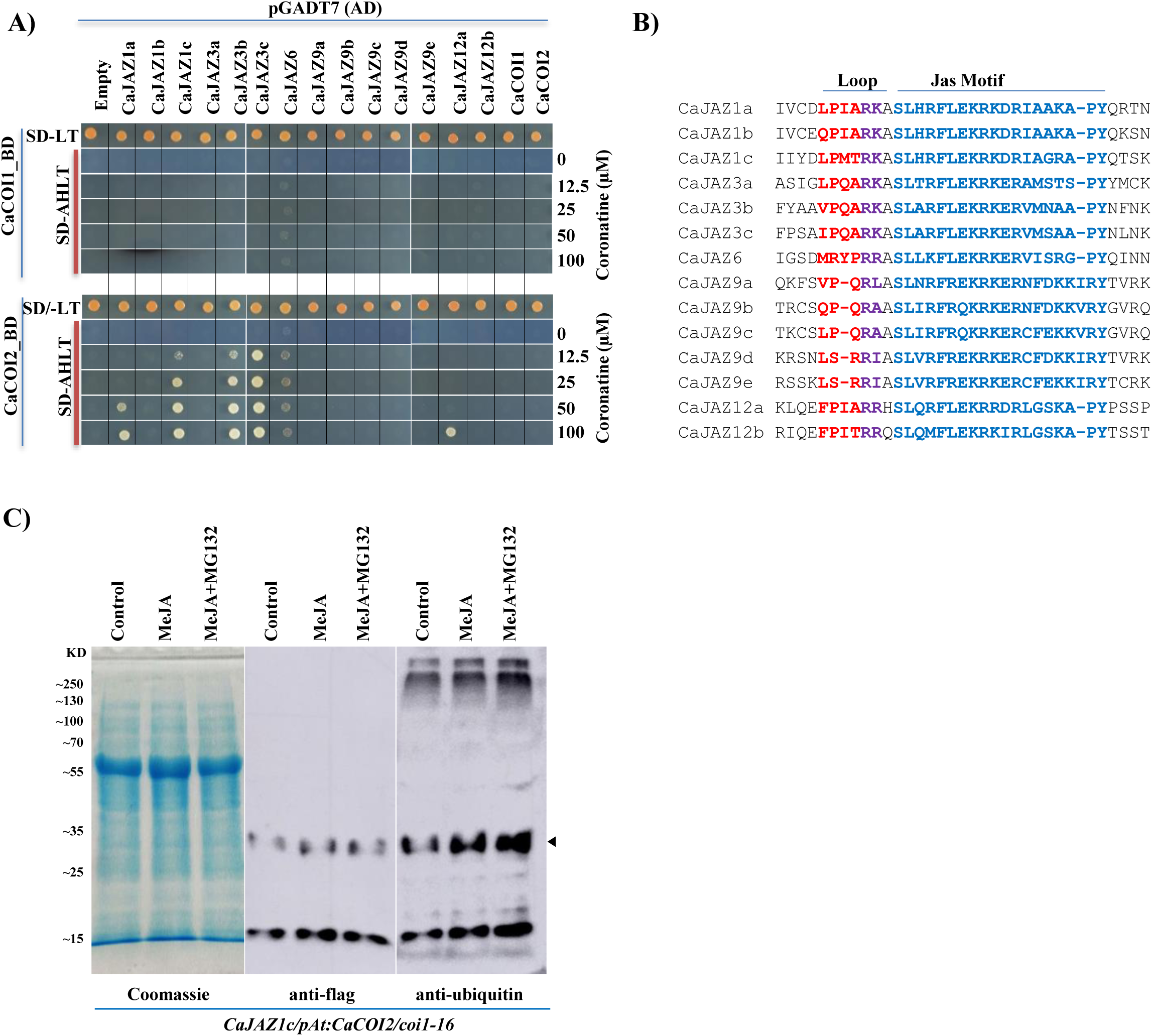
CaCOI2 facilitates ubiquitination of CaJAZ proteins in the presence of MeJA. **A)** Yeast-two-hybrid assay showing interaction of CaCOI1 and CaCOI2 with CaJAZ proteins. Upper panel in both sets represent growth of double transformants with associated genes (in pGADT7 and pGBKT7 vectors) in Y2HGold strain on SD-LT media while the lower panels represent their interaction on SD-AHLT media supplemented with indicated concentrations of coronatine. Growth was recorded at 3 days after inoculation. **B)** Protein sequence alignment of CaJAZs showing loop and Jas motif of all the identified JAZ proteins. **C)** In-vivo ubiquitination of CaJAZ1c in *pAt:CaCOI2* complemented *coi1-16* line during control, 100µM MeJA or 100µM MeJA and 50µM MG132, respectively. ∼12µg protein was loaded to each well. Coomassie staining indicates equal loading. Arrow marks the most abundant poly-ubiquitinated form of CaJAZ1c.

To unravel the degradation of JAZs by the E3 ubiquitin ligase activity of CaCOI2, we generated flag-CaJAZ1c transgenic lines in pAt:CaCOI2 complemented line (L2) in the background of *coi1-16* mutant so that the E3 ligase activity can only be imparted by the CaCOI2 but not AtCOI1 protein. The ubiquitination assay in these lines suggested higher ubiquitination of CaJAZ1c on MeJA application compared to control conditions. The addition of MG132 further enhanced the ubiquitination intensity (Figure 4C). Further, to exclude any non-specific binding of antibodies we analysed the ubiquitination pattern in the parent line i.e. *pAt:CaCOI2/coi1-16* and we observed that the bands pattern observed in *CaJAZ1c/pAt:CaCOI2/coi1-16* were specific to CaJAZ1c only (Figure S11). As these experiments were conducted in the background of *coi1-16* mutant at 22 °C, the ubiquitination of CaJAZ1c is expected to result from CaCOI2 ubiquitination activity. These results validated that CaCOI2 is a functional E3 ubiquitin ligase that participates in JA signaling and acts as JA receptor in chickpea.

### The dual localization dynamics of CaCOIs explain their differential behaviour

Next, we investigated the reason for the non-functionality of CaCOI1 despite having all known critical residues required for COI-JAZ and COI-Coronatine interactions (Figure S1). Both JAZ and COI proteins must localise to a common subcellular domain for physical interaction. Previous reports have confirmed the nuclear localization of JAZ and COI proteins (Chini et al., 2007; Thines et al., 2007; Singh et al., 2015; Yan et al., 2018). Hence, we focused on co-localization of JAZ and COI proteins. First, we predicted the subcellular localization of identified chickpea JAZ and COI proteins with Plant-mPLoc server (http://www.csbio.sjtu.edu.cn/bioinf/plant-multi/) and LOCALIZER (http://localizer.csiro.au/). Both the servers supported the nuclear localization of chickpea JAZ and COI proteins. Plant-mPLoc server supported nuclear localization of all JAZ and COI proteins while LOCALIZER supported nuclear localization of 12 JAZ proteins (except CaJAZ9b and -9c) and both COI proteins (Table S5). To confirm the nuclear localization of identified chickpea JAZ and COI proteins, we cloned CDS of all fourteen *CaJAZs* and two *CaCOIs* and generated YFP-gene fusion constructs. Using particle bombardment in onion epidermal cells, we confirmed the nuclear localization of all 13 CaJAZ proteins (Figure S12), while the 14^th^ candidate JAZ protein (CaJAZ6) was already reported localized to the nucleus in our previous study (Singh et al., 2015). To our surprise and in contrast to the previous report for Arabidopsis COI1 (Yan et al., 2018), we observed the localization of both chickpea COI proteins along the plasma membrane during normal conditions (Figure 5A-B). However, on exogenous MeJA treatment, the YFP signals was observed exclusively in the nucleus for CaCOI2, but for CaCOI1 signals were still observed along the plasma membrane and on the nuclear membrane (Figure 5A-B). To further validate this, we analysed subcellular dynamics of YFP-CaCOI2 in tobacco leaves, where we observed that CaCOI2 was associated with plasma membrane as well as the nucleus. Further, YFP signals from plasma membrane vanished within 5 minutes of MeJA treatment, making it exclusively localised to the nucleus. This observation was CaCOI2 specific as there was no impact of MeJA treatment on empty vector (YFP) fluorescence (Figure 5C-D). Considering the fast JA response and wounding caused during sample preparation, these results can well be justified. The quantitative analysis by normalizing fluorescence values in the nucleus with plasma membrane for respective time points again excluded the possibilities of technical errors like fluorescence bleaching etc. Therefore, these observations confirmed a dual localization pattern for the JA receptor where JA receptor localizes to plasma membrane/cytosol during normal conditions, but during stress i.e., in the presence of JA-Ile, they move to the nucleus to initiate JA signaling. Also, this dual localization pattern explains the non-functional behaviour of CaCOI1 by its failure to penetrate the nuclear membrane and interact with nuclear-localized JAZ proteins.

**Figure 5.**
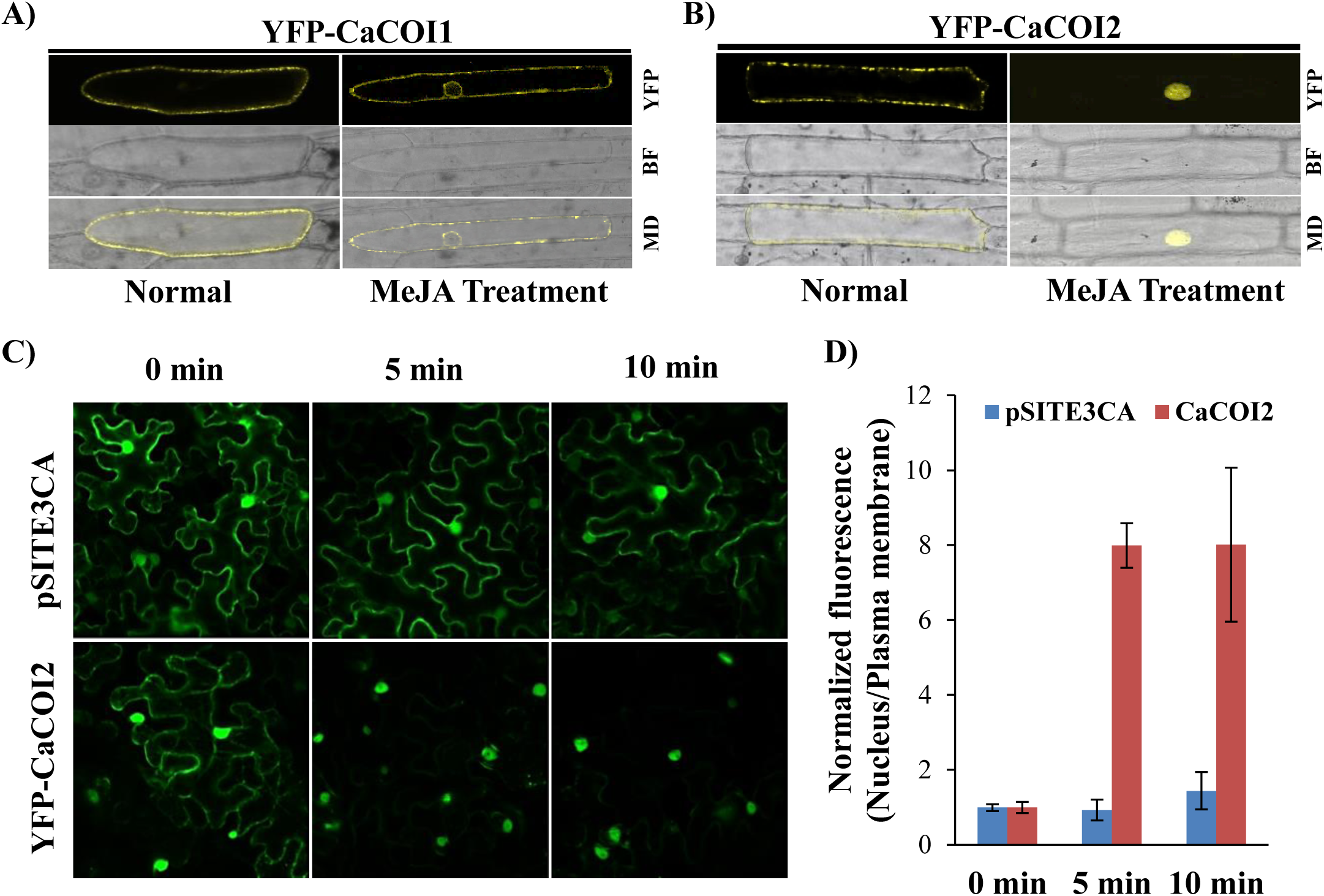
MeJA treatment promotes nuclear localisation of CaCOI proteins. **A-B)** Representative images showing subcellular localization of CaCOI1 and CaCOI2 in onion epidermal cells using particle bombardment under normal conditions and after MeJA (100 µM) treatment. For MeJA treated epidermal cells localization was analyzed after 30 min of the treatment. CaCOI1/2 were fused with YFP to produce YFP-gene fusion. YFP=Fluorescence; BF=bright field and MD=merged image. **C)** Subcellular localization of pSITE3CA (Empty vector) and YFP:CaCOI2 proteins in Nicotiana leaves. YFP signals were analysed 0 min (Control), 5 min and 10 min after 100µM MeJA treatment. **D)** Quantitative analysis of YFP signals where mean fluorescence values were first normalized to control conditions (0 min) and then average of ratio of fluorescence values of nucleus to plasma membrane was plotted with SE among the three replicates. Student’s *t-*test; \**P* < 0.05.

### LRR region of COI proteins participates in JA-mediated subcellular localization dynamics of JA receptors

To further investigate the inactive behaviour of CaCOI1, we swapped F-box region of both the CaCOI proteins with each other and analysed their interaction with CaJAZ proteins in the presence of coronatine. Interestingly, we observed that neither CaCOI1^F-box^-CaCOI2^LRR^ (F-box region of CaCOI1 fused with LRR region of CaCOI2) nor CaCOI2^F-box^-CaCOI1^LRR^ (F-box region of CaCOI2 fused with LRR region of CaCOI1) was able to interact with JAZ proteins (Figure 6A). Considering CaCOI2 has a functional F-box and LRR region, the inactive behaviour of both chimeric proteins suggested that there can be multiple additional amino acid residues distributed along F-box and LRR region, which still needed to be identified as “critical” for their functionality as a JA receptor. These results were in agreement of previous reports where multiple alleles were found associated with the AtCOI1 activity (Feys et al., 1994; Yan et al., 2009; Sheard et al., 2010; Song et al., 2021). Hence, we mapped amino acid residues of CaCOI1 and CaCOI2 with previously identified *atcoi1* alleles; however, none of these alleles explained the non-functional behaviour of CaCOI1 (Feys et al., 1994; Ellis and Turner, 2002; Yan et al., 2009; Acosta et al., 2013; Song et al., 2021, Figure S13). Interestingly, the CaCOI1^F-box^-CaCOI2^LRR^, though failed to interact with JAZ proteins, was able to localize to the nucleus on MeJA treatment, but CaCOI2^F-box^-CaCOI1^LRR^ failed to penetrate the nuclear membrane (Figure 6B). These results suggested that the LRR region of COI proteins participates in nuclear targeting.

**Figure 6.**
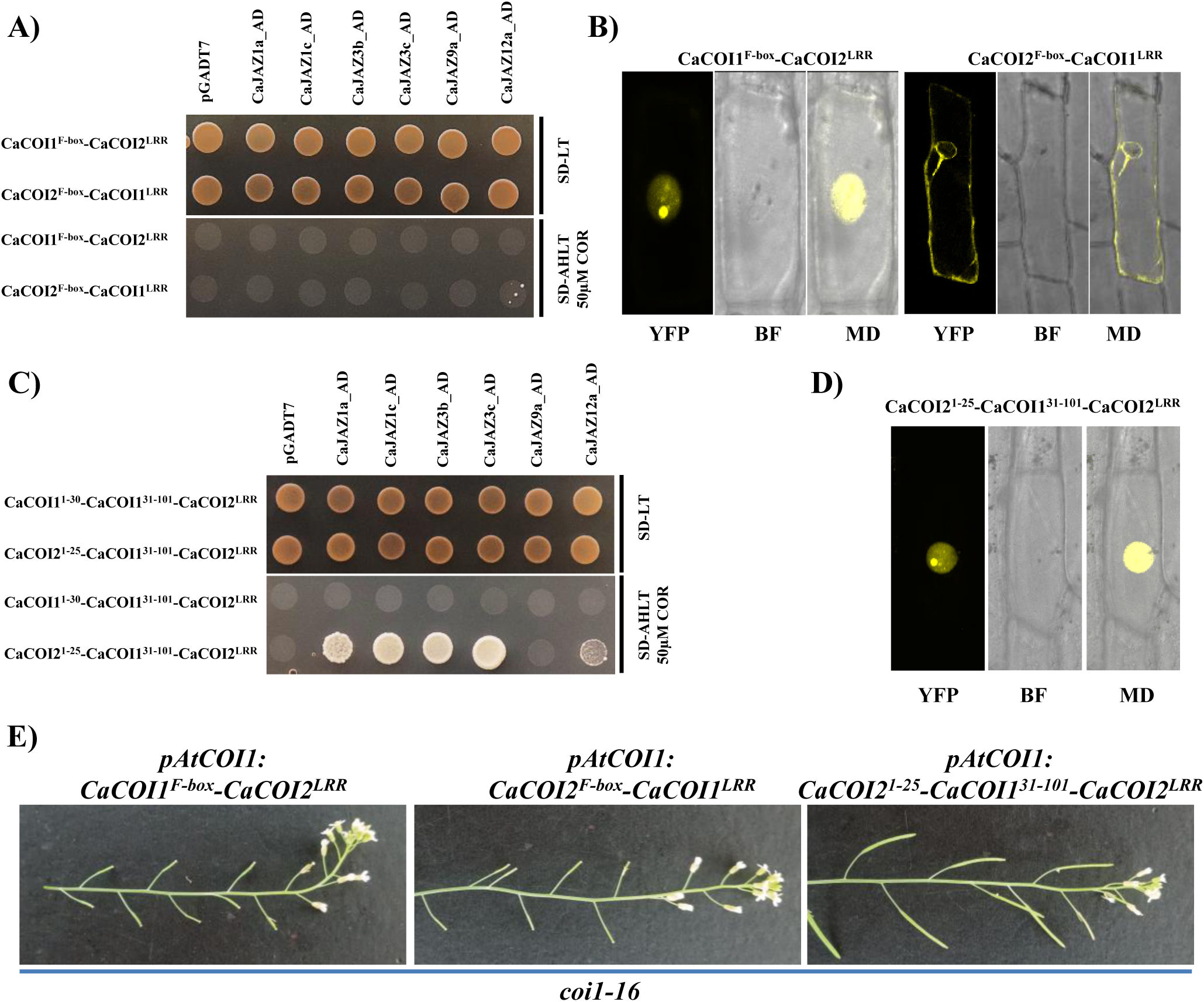
LRR region of COI proteins participates in their nuclear targeting. **A)** Yeast-two-hybrid assay showing interaction of CaCOI1^F-box^-CaCOI2^LRR^ and CaCOI2^F-box^-CaCOI1^LRR^ chimera variants of CaCOI protenis with different CaJAZ proteins. Upper panel in both sets represent growth of double transformants with associated genes (in pGADT7 and pGBKT7 vectors) in Y2HGold strain on SD-LT media while the lower panels represent their interaction on SD-AHLT media supplemented with 50µM coronatine. Growth was recorded at 3 days after inoculation. **B)** Subcellular localization of CaCOI1^F-box^-CaCOI2^LRR^ and CaCOI2^F-box^-CaCOI1^LRR^ was observed in onion epidermal cells using particle bombardment after MeJA (100 µM) treatment. **C)** Yeast-two-hybrid assay showing interaction of CaCOI1^1-30^-CaCOI1^31-101^-CaCOI2^LRR^ (i.e. CaCOI1^F-box^-CaCOI2^LRR^) and CaCOI2^1-25^-CaCOI1^31-101^-CaCOI2^LRR^ variants with different CaJAZ proteins. **D)** Subcellular localization of CaCOI2^1-25^-CaCOI1^31-101^-CaCOI2^LRR^ variant was observed in onion epidermal cells using particle bombardment after MeJA (100 µM) treatment. **E)** Representative images showing complementation of *coi1-16* mutant with different variants of chimeric JA receptors. For complementation, *coi1-16* mutant was transformed with indicated constructs (CDS driven by *AtCOI1* promoter) using Agrobacterium mediated transformation and then analyzed for silique formation at 22°C in T_1_ generation.

The chimera protein, CaCOI1^F-box^-CaCOI2^LRR^, has the functional LRR region derived from CaCOI2, indicating that the F-box domain derived from CaCOI1 might have contributed to its non-functionality. Therefore, we revisited the F-box region of COI proteins and found that a nucleophilic or acidic residue (AtCOI1^T29^, CaCOI2^D25^) remains highly conserved among functional COI proteins. However, this acidic residue is replaced with a basic residue (CaCOI1^H30^; Figure S14) in CaCOI1. Considering the highly conserved region within the F-box and the close proximity of this acidic amino acid to the AtCOI1^H54^ (a phosphate interacting residue, Sheard et al, 2010), we replaced entire N’-terminal region along with this amino acid (replacing CaCOI1^1-30^ with CaCOI2^1-25^) with F-box from CaCOI2 in CaCOI1^F-box^-CaCOI2^LRR^ resulting into a chimeric protein CaCOI2^1-25^-CaCOI1^31-101^-CaCOI2^LRR^. For this experiment, we depicted CaCOI1^F-box^-CaCOI2^LRR^ as CaCOI1^1-30^-CaCOI1^31-101^-CaCOI2^LRR^ for being the parent construct. Interestingly, CaCOI2^1-25^-CaCOI1^31-101^-CaCOI2^LRR^, which contained the conserved acidic residue (CaCOI2^D25^), was now able to interact with the JAZ proteins (Figure 6C). Also, subcellular localization confirmed the presence of CaCOI2^1-25^-CaCOI1^31-101^-CaCOI2^LRR^ in the nucleus on MeJA treatment as observed for CaCOI2 native protein and CaCOI1^F-box^-CaCOI2^LRR^ protein (Figure 6D). We further confirmed the physiological functionality of this chimeric protein by transforming *coi1-16* mutant with CDS for either CaCOI1^F-box^-CaCOI2^LRR^, CaCOI2^F-box^-CaCOI1^LRR^ or CaCOI2^1-25^-CaCOI1^31-101^-CaCOI2^LRR^ driven by AtCOI1 promoter and analysed the transformants for silique formation. Expectedly, neither of CaCOI1^F-box^-CaCOI2^LRR^ and CaCOI2^F-box^-CaCOI1^LRR^ lines were able to produce seeds, but CaCOI2^1-25^-CaCOI1^31-101^-CaCOI2^LRR^ produced siliques, thus validating our hypothesis (Figure 6E). These results confirmed that the LRR region is involved in the subcellular localization dynamics of COI proteins in response to JA perception. Further, an acidic amino acid (AtCOI1^T29^, CaCOI2^D25^) is required at this conserved position in the F-box region to facilitate interaction between COI and JAZ proteins as the subcellular localisation remained justified. We speculate that the presence of basic residue, CaCOI1^H30^, in the close proximity of highly conserved phosphate binding residue (CaCOI1^H57^ or AtCOI1^H55^ or CaCOI2^H50^) may have interfered with the phosphate binding capacity, resulting into the inactivity of the CaCOI1 protein.

### Mutations at AtCOI1^T29^ destabilizes InsP8 perception

In order to further validate our hypothesis, we analysed the structural basis of JA-Ile and InsP8 perception by AtCOI1 and if there is any change in the perception after substituting Thr29 with His29 (T29H) in AtCOI1 protein. Whereas, no significant change in the binding affinity or binding pattern was suggested w.r.t. AtCOI1/JA-Ile interaction, we observed that substituting Thr29 with His29 in AtCOI1 results into decrease in the number of hydrogen bonds (15 in AtCOI1T29 and 12 in AtCOI1H29) and even a change in orientation of ligand and participation of different residues while interacting with InsP8 (Figure 7A-B). Molecular docking analysis displays the binding energy of InsP8 to AtCOI1 as -6.42 kcal/mol, which increases to -7.21 kcal/mol when Insp8 binds to the AtCOI1/JA-Ile complex, similarly, in T29H, the binding affinity value also increases from -6.02 to -6.76 kcal/mol. Further, the docking result could indicate that Insp8 might preferentially bind to the JA-Ile bound AtCOI1. In order to further understand the dynamics of JA-Ile and InsP8 perception we performed molecular dynamics simulations. While analysing InsP8 binding to AtCOI1 ^T29^, it is evident that the protein shows a stable trajectory for RMSD while a significant decrease in Rg trajectory values indicate an increase in the compactness of the protein. However, in the AtCOI1^H29^-InsP8 complex an increase is seen in RMSD values indicating introduction of instability while an increase in the Rg trajectory values, indicates an opening of the protein fold. SASA values indicate a similar result: the AtCOI1^T29^-InsP8 complex displays a value of 281.92 nm^2^ while AtCOI1^H29^-InsP8 complex displays an increase in SASA value (286.90 nm^2^). PCA also displays an increase in Cα fluctuations of AtCOI1^H29^-InsP8 complex suggesting the inherent instability of the complex (Figure 7C-D; Figure S15). While analysing the JA-Ile binding to AtCOI1, both Rg and SASA analysis show a similar result where the ligand-bound states of T29H display a general decrease in values, indicating a decrease in the overall compactness of the protein, while the native protein upon binding with JA shows an increase in both values which is indicative of introducing fluctuations in the protein fold (Figure S15). Hence, our analysis suggests no direct involvement of Thr29 in InsP8 binding to AtCOI1 but causes a significant re-organisation resulting in destabilised AtCOI1H29-InsP8 compared to AtCOI1^T29^-InsP8 interaction. Further, JA-Ile is suggested to binds first with AtCOI1 followed by InsP8 molecule (Table S7).

**Figure 7.**
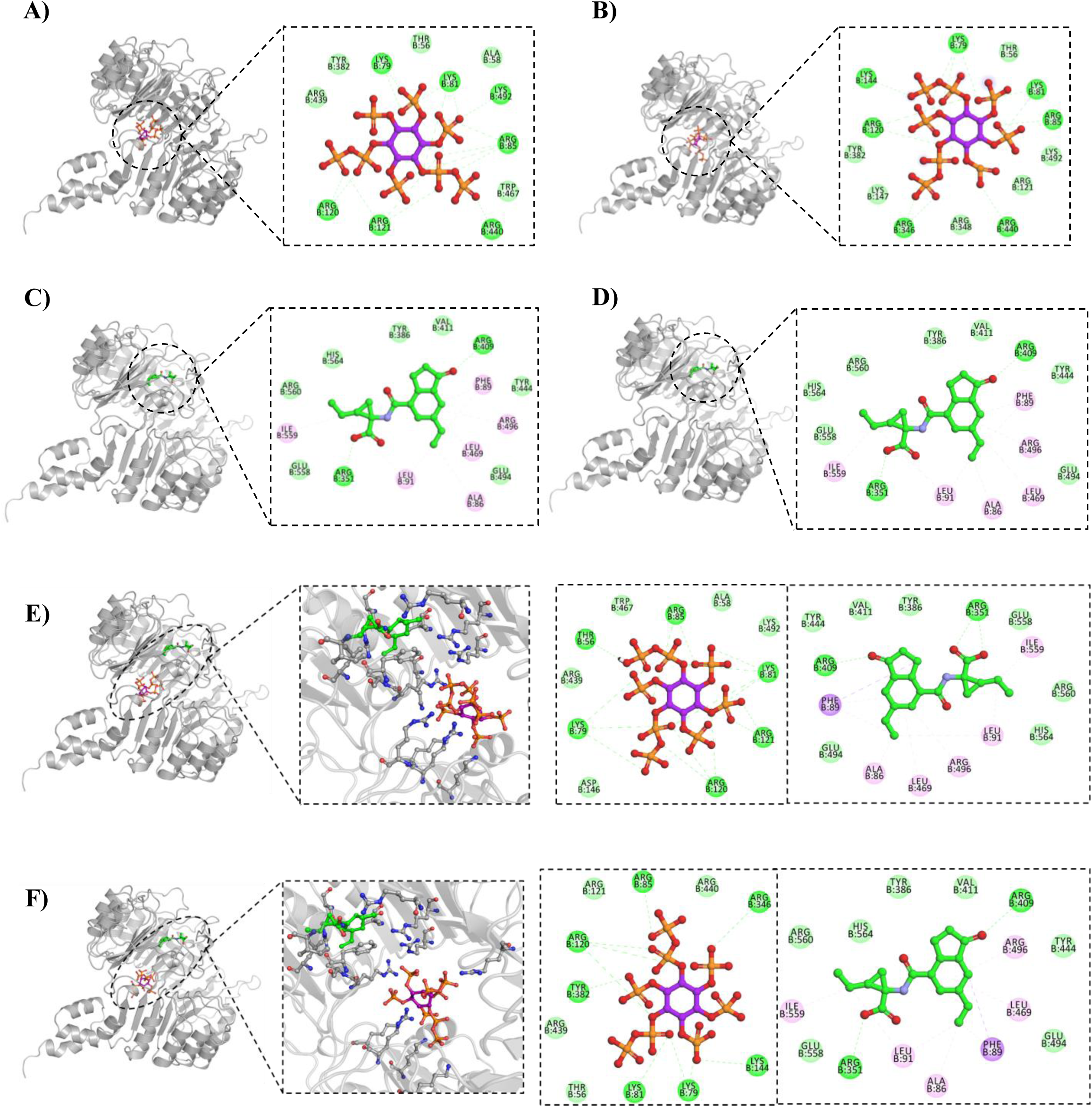
AtCOI1 F-box participates in InsP8 perception. Docking results for **(A)** AtCOI1^T29^/InsP8, **(B)** AtCOI1^H29^/InsP8, **(C)** AtCOI1^T29^/JA-Ile, **(D)** AtCOI1^T29^/JA-Ile, **(E)** AtCOI1^T29^/InsP8/JA-Ile, and **(F)** AtCOI1^H29^/InsP8/JA-Ile complex. Intermolecular interactions represented with green for hydrogen bonds, light green for Vander Waals interactions, pink and purple for hydrophobic interactions.

Therefore, these findings suggest an important role of amino acid residues at the extreme N’-terminal end of the F-box region for COI-JAZ interaction, possibly through InsP8 as an intermediate.

## Discussion

Jasmonates are known to regulate a wide range of stress responses in plants. Though JA signaling components have well been characterized in model plant Arabidopsis, the ongoing research is focused on characterizing JA signaling components in crops so that they can be utilized for improving agronomic traits. In order to bridge the missing information on JA signaling components in chickpea genome, we characterize the newly identified chickpea JA receptor COI genes (Singh et al., 2015). COI encodes for an F-box protein, a component of SCF ubiquitin ligase complex involved in protein degradation via 26S proteasome pathway (Ellis and Turner, 2002; Yan et al., 2009). In the presence of JA (JA-Ile), COI proteins interact with JA signaling repressor JAZ proteins and facilitate ubiquitination mediated degradation of JAZ proteins through 26S proteasomal pathway resulting in active JA signaling (Wasterneck and Hause, 2013). Previously, one COI homologue was identified in chickpea genome, but improved genome coverage enabled us to identify one additional potential JA receptor in chickpea (Singh et al., 2015). Interestingly, even having all the previously characterized amino acid residues in both the proteins only one of these two JA receptors was found functional (CaCOI2). So far, JA signaling components have been identified in many plants and in most of them multiple JAZ proteins are identified while only a few COI proteins characterized. Arabidopsis has a single COI protein for 12 JAZ proteins, rice has three COI for fifteen JAZ proteins and similarly, chickpea has two JA receptors against 14 JAZ proteins, and out of them only one is functional. Therefore, there must be some component determining the specificity of JA response. Our results with interaction between COI and JAZ proteins showed an increasing number of interactors with increasing concentrations of coronatine. The phenomenon seems to explain how the same signaling machinery can be so specific to a particular stress. We suspect that the ligand concentration dictates the severity and the kind of the stress and, therefore, select its interacting partners to produce a defined response. Secondly, such specificity can also be achieved by the genetic variation among these JA receptors at amino acid level. Such genetic variation has been observed in rice and Zea mays, where functional redundancy is observed in both JAZ and COI proteins (Singh et al., 2015; Qi et al., 2022). These variants of JAZ and COI proteins have differential nature for both transcriptomic and signaling aspects. In agreement of these observations, our analysis with chickpea COI proteins also suggests multiple loci participating in the functionality of CaCOI1 proteins. Our experimentation with chimeric proteins revealed that neither F-box nor LRR region of CaCOI2 was able to complement CaCOI1 revealing that genetic diversity at both F-box and LRR region need to be explored to identify all the critical amino acid residues.

The subcellular analysis of CaCOI1 and CaCOI2 revealed that COI proteins normally localize to the extra-nuclear region while they move to nucleus on JA perception. Among the two identified JA receptors, CaCOI2 was able to follow such pattern. Although CaCOI1 localization pattern was also changed on JA treatment, it failed to penetrate nuclear membrane, suggesting that the inability of CaCOI1 to penetrate the nuclear membrane rendered it a nonfunctional JA-receptor. Therefore, these results on one hand validated CaCOI2 as a functional JA-receptor, while on the other hand, suggested differential localization pattern as the basis of the non-functionality of CaCOI1 protein. Previously, AtCOI1 was found to be localized to the nucleus in root cells probably because of stress or delay during sample preparation while analyzing the signals; authors might have documented the final stage of nuclear-localized AtCOI1 protein (Yan et al., 2018). Our results with tobacco leaf infiltration validated this conclusion where we also observed CaCOI2 localized to both nucleus and plasma membrane while within 5 min of MeJA treatment most of the signal was exclusively present in the nucleus. Further, these results also suggested that JA-Ile or COR may be perceived by COI proteins prior to their interaction with JAZ proteins and that too in a different subcellular domain. After perceiving JA, CaCOI proteins move to the nucleus where they interact with JAZ proteins and degrade them to initiate JA signaling. This JA perception and signaling model fits well with the model Yan et al. suggested (2018), where AtCOI1 could not interact with JAZ1 until it was pre-treated with COR or JA-Ile. Also, COI1 crystal structure and domain architecture were found to be very close to the TIR1 (TRANSPORT INHIBITOR RESPONSE1) which is an auxin receptor and also an F-box protein containing LRRs (Tan et al., 2007). In addition to the structural similarity, TIR1 also has the InsP6 binding cavity similar to the InsP5/InsP4 in the case of COI1, suggesting a similar mode of hormonal perception among these conserved receptors. Notably, auxin perception by TIR1 model suggests that auxin first sits in the binding site and act as molecular glue for the substrates i.e., Aux/IAAs which again are transcriptional repressors and are targeted through 26S proteasomal pathway by the SCF^TIR1^ E3 ubiquitin ligase complex (Tan et al., 2007).

Based on our docking analysis, we speculated that substitution of Thr^29^ to His^29^ (T29H) may result in an unstable variant of AtCOI1 as observed in case of CaCOI1^F-box^-CaCOI2^LRR^ which remained inactive. Additionally, the docking analysis suggests JA-Ile to be perceived first by JA receptor and then by InsP8 interaction. These results in addition of subcellular localisation suggest that JA-Ile perception leads to nuclear localization of JA receptor as observed in case of both CaCOI1^F-box^-CaCOI2^LRR^ and CaCOI2^F-box^-CaCOI1^LRR^ but neither of them was able to complement the *coi1-16* mutant because of their inability to interact with JAZ proteins. These results can be explained as CaCOI1^F-box^-CaCOI2^LRR^ is inactive by virtue of T29H mutation in the F-box region which rendered it unable to perceive InsP8 while CaCOI2^F-box^-CaCOI1^LRR^ is inactive for having inactive LRR region from CaCOI1. As LRR region is known to perceive JA; therefore, JA mediated nuclear localization was not observed in the variants with inactive LRR region. Secondly, CaCOI2^1-25^-CaCOI1^31-101^-CaCOI2^LRR^ and CaCOI1^1-30^-CaCOI1^31-101^-CaCOI2^LRR^ both contained active LRR region find their way to nucleus suggesting that nuclear localization is a JA dependent phenomenon and may not be InsP8 dependent. Further, we speculate that JA is perceived first which is then followed by Insp8 as validated by our subcellular localization pattern where CaCOI1^F-box^-CaCOI2^LRR^ (which is unable to perceive InsP8) though is able to localize to nucleus, failed to interact with JAZ proteins. Therefore, to interact with JAZ proteins, COI1 first needs to interact with JA-Ile and Insp8 while both ligands are critical for active JA signaling. These observations well justified the results obtained with *atvih2* mutant, which showed lower JA signaling because of being deficient in InsP8 (Laha et al., 2015). Therefore, we propose that besides, JA-Ile content in the plant, abundance of InsP8 is also a key determinant in the initiation of JA signaling in response to any biotic and abiotic response. Interestingly, Inositol phosphates are closely related to the certain abiotic stresses like phosphate (Pi) deficiency (Ma et al., 2022). VIH2 is known to regulate InsP8 synthesis while InsP8 is proved to act as a signaling molecule driving PSR (phosphate starvation response) in Arabidopsis (Laha et al., 2015). Mechanistically, InsP8 remain elevated during phosphate-sufficient conditions acting as the glue between cytoplasmic SPX proteins and the master regulator TF i.e. PHR2. While in low Pi conditions, PHR2 can freely move to nucleus to induce PSR related gene expression because of the low abundance of InsP8 (Ma et al., 2022). Low Pi induces JA abundance and signaling in a COI1 dependent manner (Khan et al., 2016). However, it is unclear if the increase in JA content is a compensatory mechanism to maintain normal levels of JA signaling, or if downregulation of InsP8 content occurs to counteract the effects of induced JA signaling. Further research is needed to confirm the exact mechanism.

In conclusion, this study identified and characterized CaCOI2 as a functional JA receptor in chickpea. The dual subcellular dynamics of CaCOI proteins on JA treatment suggest that JA-Ile perception takes place at plasma membrane/cytosol initially by COI proteins. This perception of JA-Ile by CaCOI proteins makes them move to the nucleus, interacting with nuclear nuclear-localized JAZ proteins. Secondly, besides the JA content, InsP8 abundance also influence the severity of JA response by influencing COI-JAZs interaction as a single mutation in this phosphate interacting region destabilize the entire COI-JAZ interactions resulting into inactive COI proteins. While role of InsP8 to influence JA signaling was confirmed previously, these results extended the conclusions towards a direct role of InsP8 in JA signaling. As COI proteins are receptor for JA, which interacts with and targets multiple JAZ proteins, the allelic variations in COI proteins can be utilized to define the downstream signaling cascade in a selective way. Many alleles have already been identified for AtCOI1 protein (Figure S13) with great physiological relevance. Notably, these alleles showed differential behavior with respect to severity, signaling and growth condition (for example, *coi1-16* is a temperature sensitive mutant which regains its functionality at 16°C and *coi1-33* allele is partially male sterile). Therefore, developing/identifying alleles for selective traits would be of great agronomic importance and will further improve our understanding of JA response to generate plants better adapted to environmental stresses.

## Experimental Procedures

### *In-silico* identification of *CaCOI* and *CaJAZ* genes

CaCOIs and CaJAZs were identified in newly released chickpea databases using HMM profiles as described previously (Singh et al., 2015). The HMM profiles generated previously (for both CaCOIs and CaJAZs) were used against the newly released chickpea databases for Desi cultivar (ICC4958; https://nipgr.ac.in/CGAP3/home.php) and Kabuli (CDC frontier; https://www.icrisat.org/). Amino acid residues were aligned using ClastalX and phylogenetic analysis was done with MEGA6/ClastalX using Neighbour-Joining (N-J) method with maximum bootstrap value set to 1000. Gene structure was analysed with the GSDS server (http://gsds.gao-lab.org/) by aligning gDNA with CDS of the respective gene.

### Plant growth conditions

All experiments were done with Arabidopsis Col-gl, *coi1-16* mutant and chickpea JGK3 cultivar. Seeds of *Arabidopsis thaliana* wild-type Columbia-gl (Col-gl), *coi1-16* mutant and complementation lines of *pAtCOI1:CaCOI2* (L2 and L6) in *coi1-16* background and were sown in pot containing soilrite and agropeat in 3:1 ratio. Seeds in the pots were stratified at 4 °C in the dark condition for 3 days. Subsequently, pots were transferred to growth chamber maintained at 12 h light (at 200 µE m^−2^s^−1^ light intensity) /12 h dark conditions at temperature of 22 °C and relative humidity was maintained at 75%. Plants were bottom-irrigated with 1/4^th^ Hoagland nutrient solution and water alternatively. Floral-dip-based transformation for complementation of *coi1-16* mutant (CS67817; TAIR) was done at 16 ℃ and 16h/8h photoperiod with growth mentioned above. Mutation in *coi1-16* was validated as described (Adams and Turner, 2010; Table S1).

### Complementation of *coi1-16* mutant

To complement *coi1-16* with chickpea *COI* genes, 2X35S promoter in Gateway^TM^ compatible pMDC32 binary vector was replaced with 1056 bp of *AtCOI1* promoter (*pAtCOI1)* from Col-gl. After that, CDS of CaCOI1 and CaCOI2 from JGK3 genotype were cloned downstream of *pAtCOI1* in pMDC32 vector. Using these constructs, multiple transgenic lines were generated following *Agrobacterium*-mediated transformation (Clough and Bent, 1998).

### KI staining of seeds

In order to visualize seeds within siliques of Arabidopsis, representative siliques were taken and immersed in 2 ml of 1% (w/v) Iodine-Potassium Iodide (KI-I_2_) solution for approximately 12hrs for starch staining and then kept in 100% ethanol for chlorophyll removal till clear siliques were visible. Staining and destaining processes were done in dark conditions with shaking and photographs were captured with stereo-zoom microscope.

### Gene expression analysis

One-month-old Arabidopsis plants of Col-gl, *coi1-16* and complemented lines (grown at 22°C) were sprayed with either DMSO or 100uM MeJA and then leaf tissues were collected on different time points including 0 min (control), 30 min, 1h and 2h after Me-JA spray. Total RNA extraction, cDNA synthesis and expression analysis were done as described previously (Singh et al., 2015). Expression analysis was done using qRT-PCR with Arabidopsis gene-specific primers from coding region. Primers were designed using PRIMER EXPRESS version 3.0 (PE Applied Biosystems™, USA). Primer pairs were cross-checked for their specificity using NCBI primer blast tool. qRT-PCR was performed with Fast SYBR® Green Master Mix and relative gene expression analysis was done using the ΔΔCt method considering *AtACT2* as internal control (Table S1).

### Bacterial pathogens and inoculum preparation

The bacterial pathogen, *Pseudomonas syringae* pv. tomato DC3000 (*Pst* DC3000), either with or without the pDSK-GFPuv plasmid (Wang et al., 2007), was cultured in King’s B (KB) medium (liquid) with antibiotics at 28 °C with constant shaking at 150 rpm. The bacterial culture was grown overnight until the optical density at 600 nm (OD600) reached 0.4. Subsequently, cells were harvested by centrifugation at 4000 rpm for 10 minutes, followed by thorough washing in sterile water. This washing process was repeated three times, and the cells were then re-suspended to the desired concentration in sterile water.

### Plant inoculation*, in-planta* bacterial multiplication assay and disease index analysis

Pathogen infection experiment was done with 30-day-old plants. Bacterial suspension of *Pst* DC3000 at concentration of 5 × 10^5^ colony-forming units (CFU)/mL was used for syringe-infiltration on the abaxial surface of leaves using a needleless syringe. For mock treatment, sterile water was used to infiltrate the leaves of plants. After inoculations, plants were kept in growth chamber at 20°C. Bacterial multiplication assay was done by collecting the leaf samples at 0, 1, 2, and 3 dpi. The bacterial multiplication numbers were quantified by completely homogenizing the leaf discs of 0.5 cm^2^ area in sterile water. The serial dilutions were made and the suitable dilution was then plated on KB agar medium plate. Bacterial multiplication number was estimated using the following formula (Wang et al., 2007):

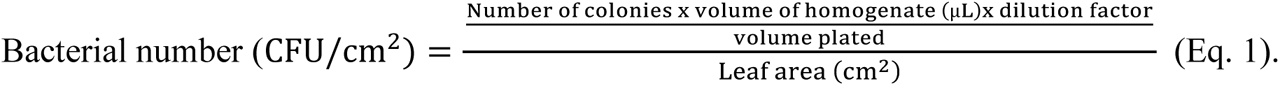

The chlorotic and necrotic disease symptoms were scored in pathogen-infected plants at 3 dpi and the disease index was calculated.

### *In-planta* imaging of bacterial population

The bacterial population of *Pst* DC3000 expressing pDSK-GFPuv plasmid was observed at cellular level in the apoplastic spaces using a confocal microscope (Leica TCS SP8, Leica Microsystems, Wetzlar, Germany). Leaf samples were harvested at 2 dpi from the plants infiltrated with GFPuv labelled bacteria. Bacterial population imaging was done in these leaf samples using 63x water immersion objective with excitation at 488 nm and emission at 550 nm.

### Interactions among CaJAZ and CaCOI proteins using Y2H assay

To analyse the interactions among CaJAZ proteins and with CaCOI proteins, CDS region of each gene was amplified from JGK3 cultivar using primer sets mentioned in table S1 and cloned in pGADT7 and pGBKT7 vectors. Subsequently, the indicated combinations of pGADT7 and pGBKT7 clones were co-transformed in Y2HGold yeast strain and transformants were selected on SD-LT media. Possible interactions among the candidate proteins was analysed on SD-HLT and SD-AHLT. For analysing Interactions between CaJAZ and CaCOI proteins SD-AHLT media was supplemented with indicated concentration of coronatine. Coronatine was first dissolved in DMSO which was then diluted with SD-AHLT media to the final stated concentration. For equal loading OD_600_ = 0.5 was used for each clone.

### Subcellular localization of CaJAZs and CaCOIs proteins

YFP-fusion constructs were generated for CaJAZ and CaCOI proteins using their CDS cloned in Gateway^TM^ compatible binary vector. These resultant YFP-gene reporter constructs were then transformed in onion epidermal cells using particle bombardment method (Singh et al., 2015). The transformed onion epidermal cells were inoculated on 1/2MS agar plates for ∼12h in the dark and then visualized under a confocal microscope (Leica SP8) for YFP signals. The transformed cells were visualized for the subcellular localization of CaCOI proteins for YFP signals. For treating onion epidermal cells with MeJA, cells were bathed with 100µM of MeJA before visualising the YFP signals. For analysing subcellular dynamics in Nicotiana leaves, YFP (pSITE-3CA) and YFP-CaCOI2 were infiltrated in tobacco leaves as per Li et al., 2008. For MeJA treatment, leaf discs were bathed in 100µM MeJA and samples were visualised under confocal microscope (ZEISS LSM 880) after 5 min and 10 min after the treatment. ImageJ was used to quantify mean fluorescence signals in the nucleus and plasma membrane by selecting a constant area within the image. Graphs were plotted with the average mean fluorescence values normalized to the control conditions after dividing the fluorescence in the nucleus by plasma membrane.

### Phenotypic analysis

For MeJA mediated root growth inhibition, seven days old seedlings grown on 1/2MS media were transferred to either 1/2MS supplemented with DMSO (control) or 100µM MeJA for next seven days. After the treatment, images were captured and primary root and lateral root length was analysed using ImageJ software (https://imagej.nih.gov/ij/). Percentage change was calculated w.r.t. the control conditions of the same construct. Anthocyanin accumulation and root hair imaging were done with stereo-zoom microscope. For nutrient deficiency experiments, homogenously grown seedlings of Col-gl, *coi1-16* and pAt:CaCOI2/*coi1-16* (L2) were transferred to either ½ MS media or ½ MS media with no external source of phosphate. Growth was analysed after 5 days of the treatment.

### Protein isolation and immunoblot analysis

In order to confirm the E3 ligase activity of CaCOI2, and CaJAZs as its targets, flag tagged CaJAZ1c was overexpressed in *coi1-16* mutant complemented with CaCOI2 (pAtCaCOI2/L2). Experiments were performed with homozygous lines. Briefly, transgenic lines homozygous for both CaJAZ1c and CaCOI2 in *coi1-16* mutant were grown at ½ MS-agar media for 15 days and then transferred to liquid ½ MS media containing DMSO (Control), 100µM MeJA and 100µM MeJA + 50µM MG132 for 3 h followed by snap freezing with liquid nitrogen. Total protein was isolated using protein extraction buffer containing [50mM phosphate buffer (pH 7.6), 5mM MgCl_2_, 100mM NaCl, 0.05% T-20 and protease inhibitor cocktail (Takara)]. Whole seedlings were homogenised in extraction buffer followed by incubation on ice for 30 min. The homogenate was centrifuged at 13000 rpm at 4 °C for 15 min and supernatant was collected.

∼10-12µg of total protein was loaded to SDS-PAGE for immunoblot analysis. After transferring to PVDF membrane, the membrane was blotted with anti-flag (Invitrogen #740001) or anti-ubiquitin antibodies (Invitrogen #PA1-10023) and detected with HRP conjugated IgG (Invitrogen #31460) using Clarity^TM^ western ECL detection system (Biorad #170-5061).

### Molecular docking and dynamics simulations

As the only available crystal structure of AtCOI1 (3OGM) lacks loop C region (Sheard et al., 2010), we first modelled the complete structure of the protein using SWISS-MODEL (Waterhouse et al., 2018), ModWeb [Pieper et al., 2014], and AlphaFold web (Jumper et al., 2021) which were energy minimized via YASARA (Land and Humble, 2018) and GalaxyRefine (Heo et al., 2013). All the models were evaluated and the best suitable model (AtCOI1 from SWISS MODEL, minimized with GalaxyRefine) was selected for molecular docking analysis (Table S6). The derived model was mutated to T29H through mutagenesis plugin in PyMoL. The ligand (JA and InsP8), T29H model and AtCOI1 model were prepared for docking through MGLTools 1.5.6. A site directed docking approach was utilized in AutoDock v4.2 which was performed with the Lamarckian Genetic Algorithm with 1000 runs, a population size of 300, and the maximum number of generations and evaluations were 27,000 and 25,00,000, respectively (Morris et al., 2009). Each ligand was docked to their experimentally hypothesized locations in both native AtCOI1 and T29H. Furthermore, to predict any preferential binding between both ligands, the AtCOI1-T29H-Jile complexes were docked with Insp8 molecules and vice versa. The docking results were analysed based on the obtained clusters, binding energy, intermolecular interactions, and docking pose, after which the best orientation was utilized for molecular dynamics (MD) simulations. MD simulation were performed via GROMACS by employing the CHARMM forcefield in a TIP4P solvated system neutralized with appropriate counter ions (Van Der Spoel et al., 2005). The whole system was energy minimized using the steepest descent model method to a final value of 100 kJ/mol/nm, and subsequently, the system was equilibrated for temperature (373 K) and pressure (1 bar) for 1 ns. Finally, a production MD was run for 100 ns, which performs 50 million steps with a time interval of 2 fs. The results for MD were analysed using several parameters, such as Root mean square deviation (RMSD), Root mean square fluctuation (RMSF), Solvent accessible surface area (SASA), Radius of gyration (Rg) and, Principal component analysis (PCA).

## Acknowledgements

A part of this research is funded by the MK Bhan Young Researcher Fellowship, DBT, India to APS and JG was supported by the NIPGR core grant and Swarnajayanti fellowship (SB/SJF/2019-20/07) by DSR-SERB, India. We acknowledge NIPGR and NABI confocal microscopy facilities.

## Authors Contribution

APS conceptualised, designed and performed most experiments and wrote manuscript with the help from co-authors. MS-K and UF contributed pathogen infection assays data. AKS and ES did computational analysis and added relevant text. JG conceptualised the project, supervised the study and wrote MS.

## Supporting Information

**Figure S1.** Amino acid sequence alignment for AtCOI1 and identified CaCOIs.

**Figure S2.** Identified chickpea *COI* homologues are similar to Arabidopsis *COI1*.

**Figure S3.** Comparison of CDS (A) and amino acid (B) sequence of identified *CaCOI* genes/proteins in CDC frontier and JGK3 varieties of Kabuli chickpea.

**Figure S4.** Complementation strategy for *coi1-16* Arabidopsis mutant using CaCOIs.

**Figure S5.** *CaCOI2* restores JA sensitivity and signaling in *coi1-16* mutant.

**Figure S6.** *CaCOI2* restores sensitivity against phosphate deficiency to *coi1-16* mutant.

**Figure S7.** Nomenclature of JAZ homologues in chickpea.

**Figure S8.** Comparison of CDS sequence of identified *CaJAZ* genes in CDC frontier and JGK3 varieties of Kabuli chickpea.

**Figure S9.** CaJAZ proteins interacts with each other to form homo and hetero-dimers.

**Figure S10.** CaCOI proteins fail to interact with CaJAZ proteins in absence of coronatine.

**Figure S11.** In-vivo ubiquitination pattern in *pAt:CaCOI2* complemented *coi1-16* lines.

**Figure S12.** Subcellular localization of identified JAZ proteins.

**Figure S13.** Multiple alleles are associated with the functionality of COI1 as JA receptor

**Figure S14.** Amino acid sequence alignment showing conserved acidic amino acid at AtCOI1^T29^ position.

**Figure S15.** Docking analysis of AtCOI1 with InsP8 and JA-Ile.

**Table S1.** List of primers used in the study.

**Table S2.** Complementation frequency of transformation events.

**Table S3.** Summary of JA associated genes identified in different chickpea genome databases.

**Table S4:** CaJAZ proteins interacts with each other to form homo and hetero-dimers.

**Table S5.** Predicted subcellular localization of identified JA associated genes.

**Table S6.** Evaluation of homology modelling derived models of AtCOI1.

**Table S7:** Docking interactions between AtCOI1, T29H and respective ligands (JA-Ile and InsP8).

