## Supplementary material for "Dual localization of JA receptor, CaCOI2, explains JA perception dynamics in chickpea"

```
                F-box
AtCOI1   M-EDPDIKRCKLSCVATVDDVIEQVMTYITDPKDRDSASLVCRRWFKIDSETREHVTMAL   59
CaCOI1   MEEKAACGGVGRTSTKLSDVVLDCVMPYIHDPKDRDAVSQVCRRWYELDSLTRKHVTIAL   60
CaCOI2   MTEDRSVKTNRVV-----DVVLDCVIPYIDDPKDRDAVSQVCRRWYELDSLTRKHVTIAL   55

                                           LRR starts
AtCOI1   CYTATPDRLSRRFPNLRSLKLKGKPRAAMFNLIPENWGGYVTPWVTEISNNLRQLKSVHF   119
CaCOI1   CYTTTPDRLRRRFPHLESLKLKGKPRAAMFNLIPENWGGFVTPWVKEISKYFDCLKSLHF   120
CaCOI2   CYTTTPSRLRRRFPHLESLKLKGKPRAAMFNLIPEDWGGHVTPWIKEISHYFDCLKSLHF   115

AtCOI1   RRMIVSDLDLDRLAKARADDLETLKLDKCSGFTTDGLLSIVTHCRKIKTLLMEESSFSEK   179
CaCOI1   RRMIVTDSDLQILARSRCNSLHALKLEKCSGFSTDGLYYVGRFCKNLRVLFMEESSVVEK   180
CaCOI2   RRMIIQDSDLKLLARSRGHVLQSLKLDKCSGFSTHGLRFIGRFCRSLKVLLLEESTIVEN   175

AtCOI1   DGKWLHELAQHNTSLEVLNFYMTEFAKISPKDLETIARNCRSLVSVKVGDFEILELVGFF   239
CaCOI1   DGEWLHVLALNNTVLETLNFYLTDIANVRIQDPELIAKNCPNLVSVKITDCEILNLMNFF   240
CaCOI2   DGNWLHELALNNTVLEFLNFYLTDIVDVKVQDLELLAKNCPNLVSVKITDCEILDLVNFF   235

AtCOI1   KAAANLEEFCGGSLNEDIGMPEKYMNLVFPRKLCRLGLSYMGPNEMPILFPFAAQIRKLD   299
CaCOI1   RYASSLEEFCGGSYNED---PEKYSAISLPAKLSRLGLTYIGKNEMPFVFPYAAMLKKLD   297
CaCOI2   RNATALEEFCGGTYNEE---PERYSSVSLPAKLCRLGLTYIGKNELPIVFMYAAALKKLD   292

AtCOI1   LLYALLETEDHCTLIQKCPNLEVLETRNVIGDRGLEVLAQYCKQLKRLRIERGADEQGME   359
CaCOI1   LLYAMLDTEDHCTLIQKCPNLEVLESRNVIGDRGLEVLASCCKKLRRLRIERGDDDQGME   357
CaCOI2   LLYAMLDTEDHCMLFQKCPNLEVLETRNVIGDRGLEVLGHCCKRLKRLRIERGDDDQGME   352

AtCOI1   DEEGLVSQRGLIALAQGCQELEYMAVYVSDITNESLESIGTYLKNLCDFRLVLLDREERI   419
CaCOI1   DEEGIVSQRGLIALSQGCPELEYMAVYVSDITNASLEHIGTHLKNLCDFRLVLLDREEKI   417
CaCOI2   DEEGTVSHRGLIALSQGCTELEYLAVYVSDITNASLEQIGTHLKNLCDFRLVLLDHEEKI   412

AtCOI1   TDLPLDNGVRSLLIGCKKLRRFAFYLRQGGLTDLGLSYIGQYSPNVRWMLLGYVGESDEG   479
CaCOI1   SDLPLDNGVRALLRGCDKLRRFALYLRPGGITDVGLGYIGQYSPNVRWMLLGYVGETDAG   477
CaCOI2   SDLPLDNGVRALLRGCDKLKRFALYLRRGGLTDIGLGYIGQYSQNVRWMLLGYVGETDAG   472

AtCOI1   LMEFSRGCPNLQKLEMRGCCF-SERAIAAAVTKLPSLRYLWVQGYRASMTGQDLMQMARP   538
CaCOI1   LLEFSKGCPSLQKLEMRGCSFFSEYALAIAATRLTSLRYLWVQGYGASPSGRDLLAMARP   537
CaCOI2   LLEFAKGCPSLQKLEMRGCSFFSEHALAVAATQLTSLRYLWVQGYGASPSGRDLLAMARP   532

AtCOI1   YWNIELIPSRRVPEVNQ-QGEIREMEHPAHILAYYSLAGQRTDCPTTVRVLKEPI-----   592
CaCOI1   YWNIELIPSRRVVVKNQ-QDELVAVEHPAHILAYYSLAGPRSDFPDTVIPLDPAAYY---   593
CaCOI2   FWNIELIPSRQVAISNNMGEPLVVVEHPAHILAYYSLAGQRSDFPDTVVPLNPATYVNAY   592

AtCOI1   ---   592
CaCOI1   ---   593
CaCOI2   SCV   595
```

● Phosphate Interacting residues ● JAZ1 degron loop contacting residues
● JA-Ile/COR contacting residues ● JAZ1 degron helix contacting residues

**Figure S1. Amino acid sequence alignment for AtCOI1 and identified CaCOIs.** Protein sequence alignment showing conserved residues among the three COI homologues. AtCOI1 and CaCOIs amino acid residues were aligned using the ClustalX program. (Sheard et al., 2010; Lee et al., 2013; Qi et al., 2022).

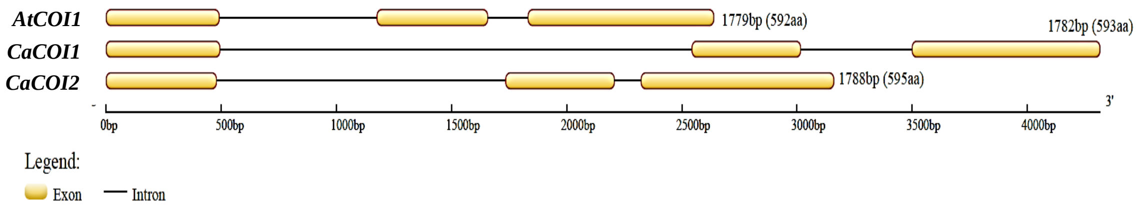


**Figure S2. Identified chickpea *COI* homologues are similar to Arabidopsis *COI1*.** Exon intron structure of *AtCOI1*, *CaCOI1* and *CaCOI2* genes. gDNA and CDS was aligned using GSDS server (http://gsds.gao-lab.org/).

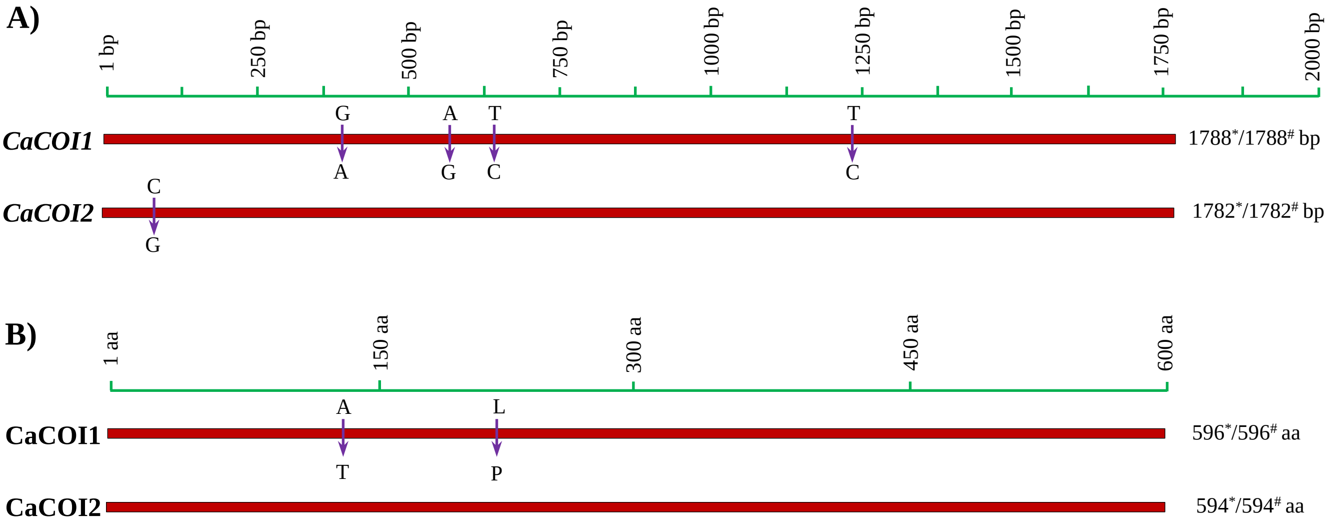


**Figure S3. Comparison of CDS (A) and amino acid (B) sequence of identified *CaCOI* genes/proetins in CDC frontier and JGK3 varieties of Kabuli chickpea.** *CDS/aa length as observed in JGK3 cultivar, #CDS/aa length as provided for CDC frontier by Varshney et al. (2013). Red boxes indicate the CDS/aa length of identified genes, while arrow indicates SNP change from CDC frontier to JGK3 cultivar.

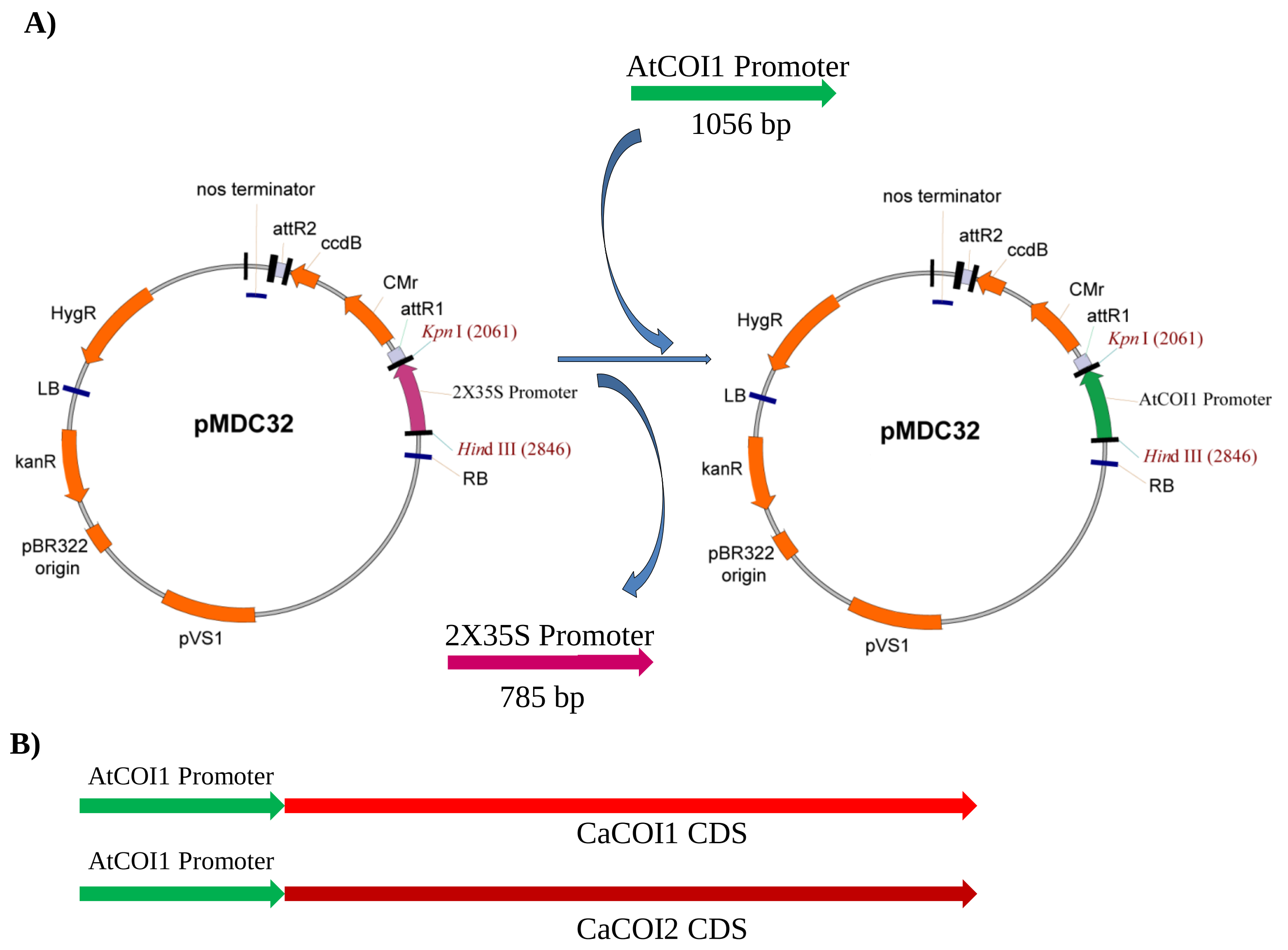


**Figure S4. Complementation strategy for *coi1-16* Arabidopsis mutant using CaCOIs.** A) Integration of *AtCOI1* promoter in pMDC32 vector. A 1056 bp *AtCOI1* promoter *(pAtCOI1)* was amplified from Col-gl gDNA and 2X35S promoter was replaced with *pAtCOI1* using KpnI and HindIII restriction sites. B) Schematic diagram showing constructs of CaCOI1 or CaCOI2 CDS being driven by *pAtCOI1* for complementation assays. After integration of pAtCOI1 in pMDC32 vectors, the resultant vector was used as GATEWAY$^{TM}$ compatible destination vector for cloning of *CaCOI1* and *CaCOI2* CDS under AtCOI1 promoter.

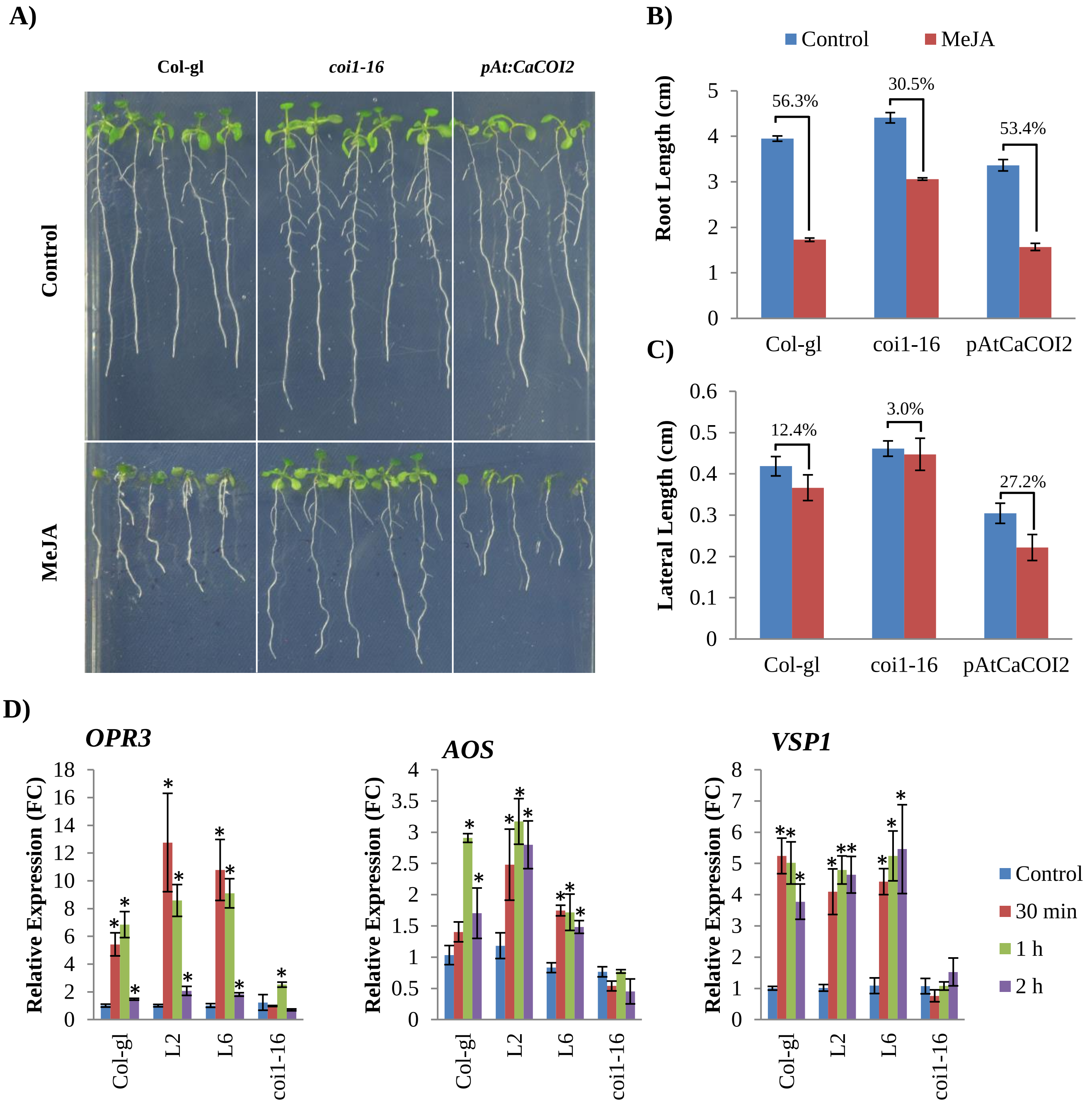


**Figure S5. *CaCOI2* restores JA sensitivity and signaling in *coi1-16* mutant. A)** Representative images showing growth pattern of Col-gl, *coi1-16* mutant and *CaCOI2* complemented line of *coi1-16* mutant *pAt:CaCOI2* (L2) under normal and 100µM MeJA treatment. **B)** Quantitative analysis of primary root length of Col-gl, *coi1-16* and *pAt:CaCOI2* (L2) complemented line under normal and 100µM MeJA. Each bar represents average of at least 20 replicates with SE among the replicates. **C)** Quantitative analysis of lateral root length of Col-gl, *coi1-16* mutant and *pAt:CaCOI2* (L2) complemented line under normal and 100µM MeJA. Each bar represents average of at least 11 replicates with SE among the replicates. **D)** Expression profiling of *OPR3*, *AOS* and *VSP1* gene in Col-gl, *pAt:CaCOI2* (L2, L6) and *coi1-16* mutant under control and 100µM MeJA treatment. Samples were collected before (Control) and after 30min, 1 h and 2 h of MeJA foliar spray. Each bar represents average of at least three biological replicates with SE among the replicates. * shows the significance change w.r.t. control conditions, Student's *t*-test.

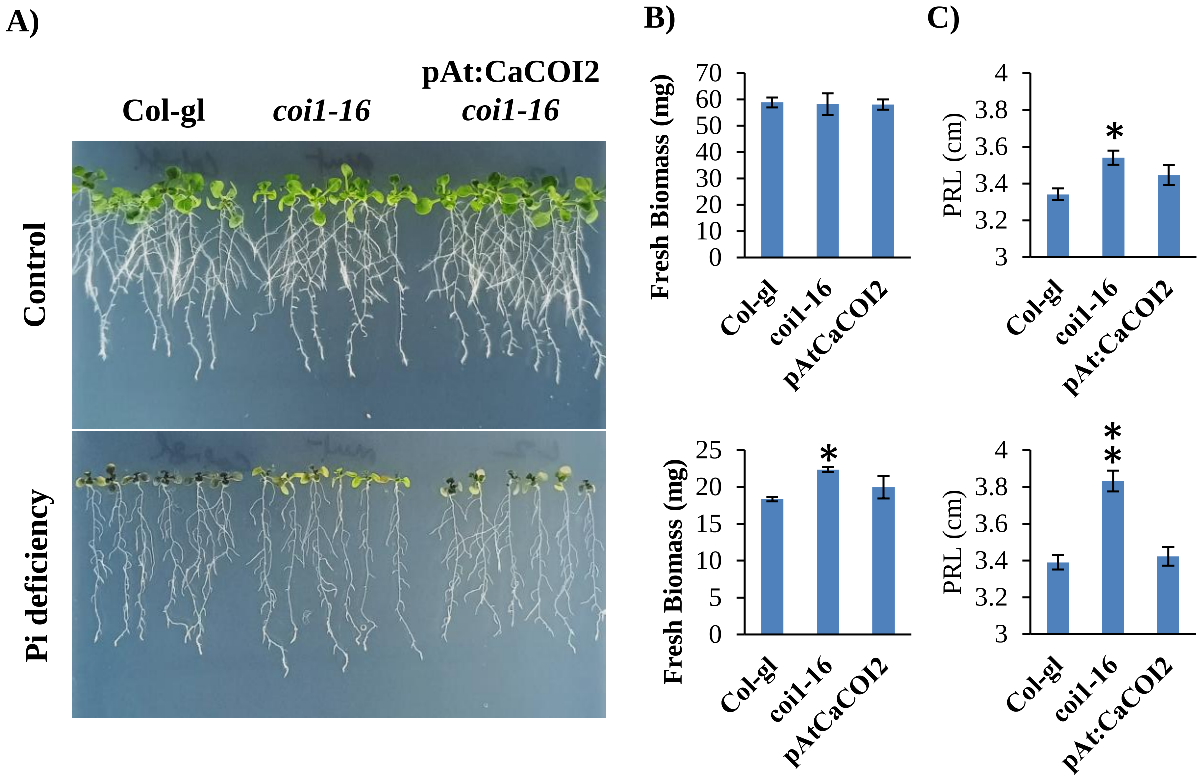


**Figure S6. *CaCOI2* restores sensitivity against phosphate deficiency to *coi1-16* mutant. A)** Representative images showing growth pattern of Col-gl, *coi1-16* mutant and *CaCOI2* complemented line of *coi1-16* mutant *pAt:CaCOI2* (L2) under normal and phosphate deficiency treatment. **B)** Quantitative analysis of fresh weight (B) and primary root length (C) of Col-gl, *coi1-16* and *pAt:CaCOI2* (L2) complemented line under normal and phosphate deficient conditions. Each bar represents average of at least 38 replicates with SE among the replicates. For fresh weight six seedlings were pooled to consider as 1 replicate and average of 8 replicates was plotted with SE among the replicates. * shows the significance change w.r.t. control conditions, Student's *t*-test.

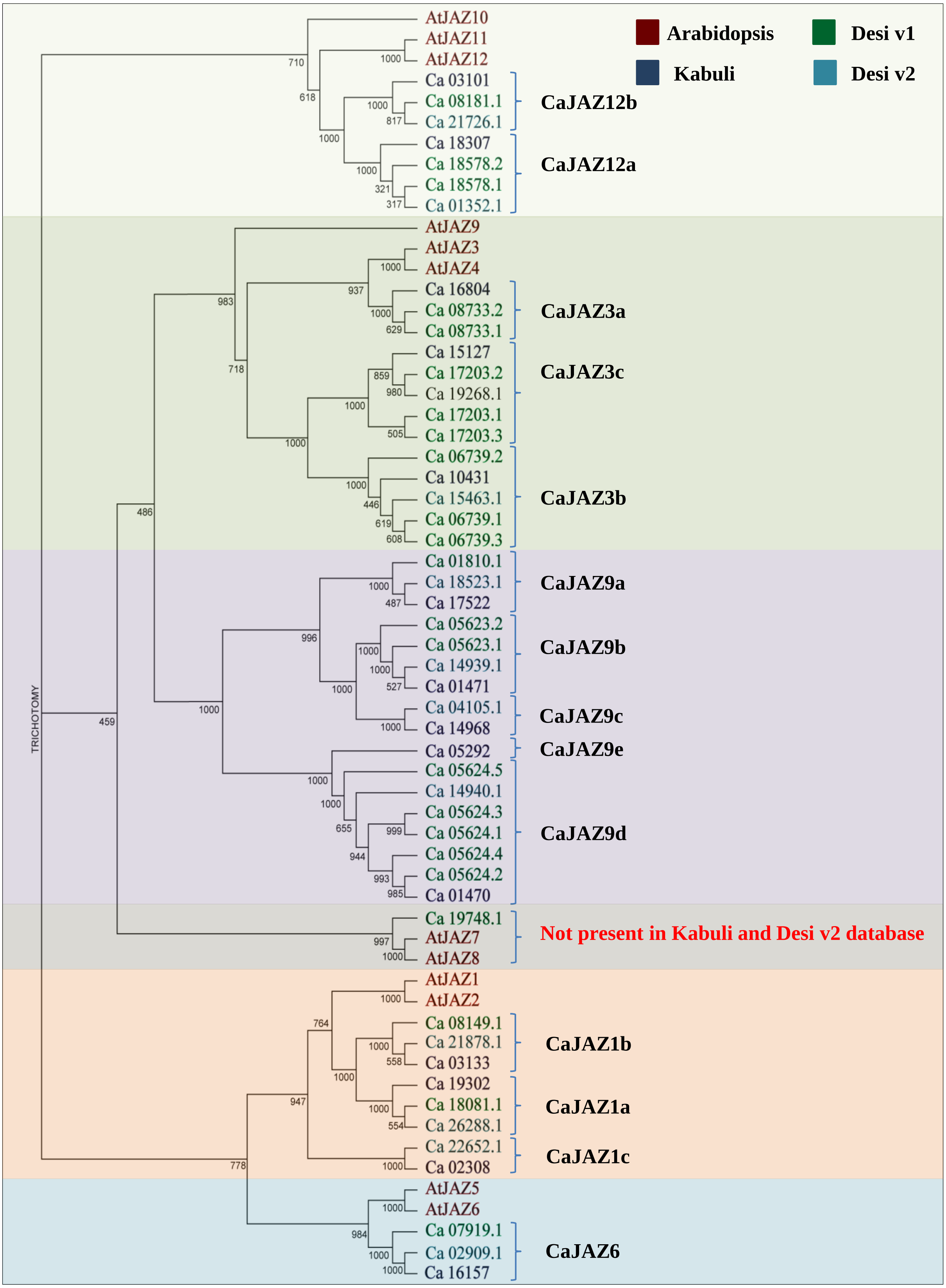


**Figure S7. Nomenclature of JAZ homologues in chickpea.** In order to name the identified JAZ homologues in chickpea, amino acid sequences of JAZ proteins were retrieved from released Kabuli, Desi [Version 1 (v1) and Version 2 (v2)] chickpea and Arabidopsis genome databases. Peptide sequences were aligned using ClustalX and phylogenetic tree was generated using N-J method with maximum bootstrap value =1000.

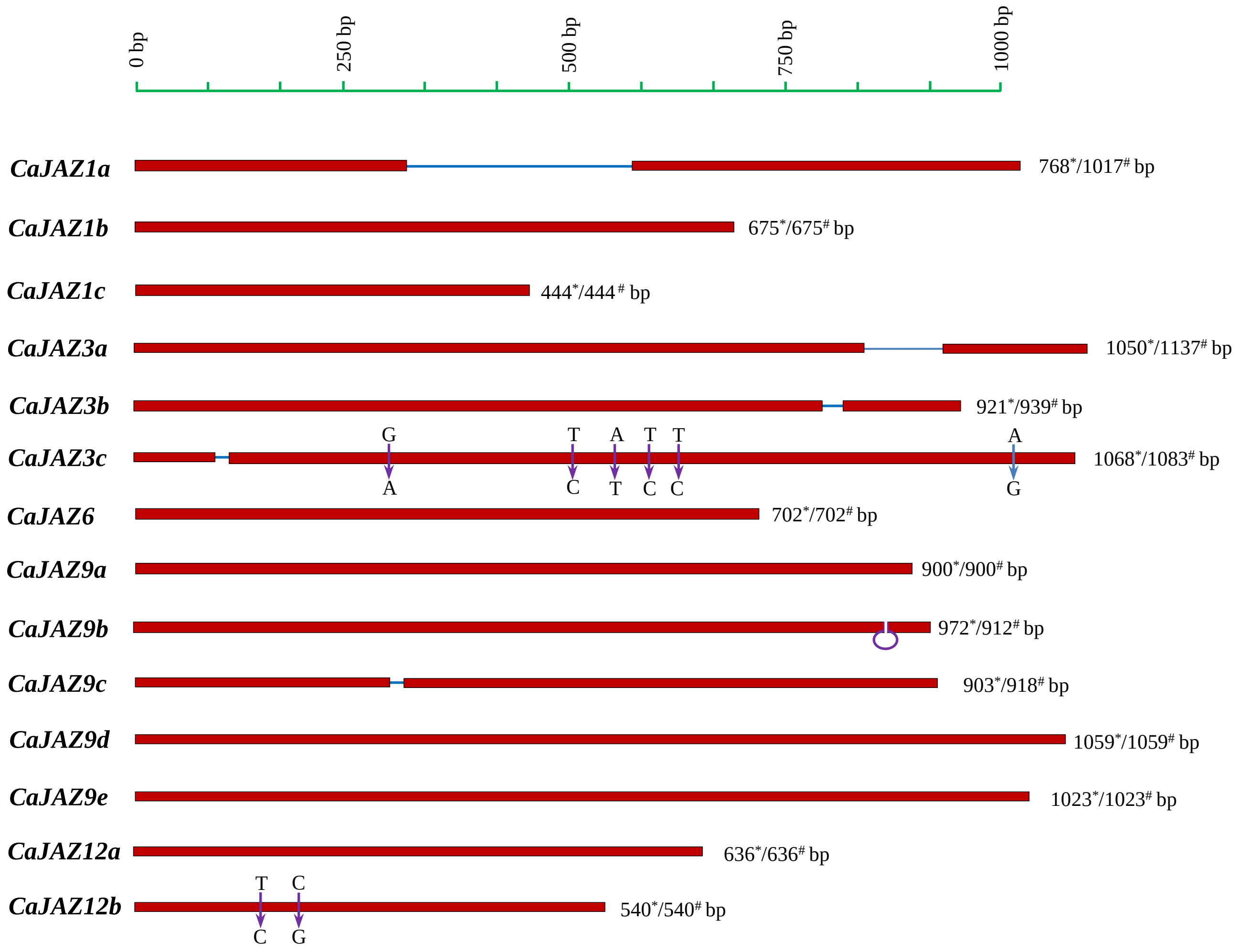


**Figure S8. Comparison of CDS sequence of identified *CaJAZ* genes in CDC frontier and JGK3 varieties of Kabuli chickpea.** Comparison of CDS sequence of identified *CaJAZ* genes. *CDS length as observed in JGK3 cultivar, #CDS length as provided for CDC frontier by Varshney et al. (2013). Red boxes indicate the CDS length of identified genes, blue line indicates the deletion, loop indicates insertion in JGK3 cultivar as compared to CDC frontier, while arrow indicates SNP change from CDC frontier to JGK3 cultivar.

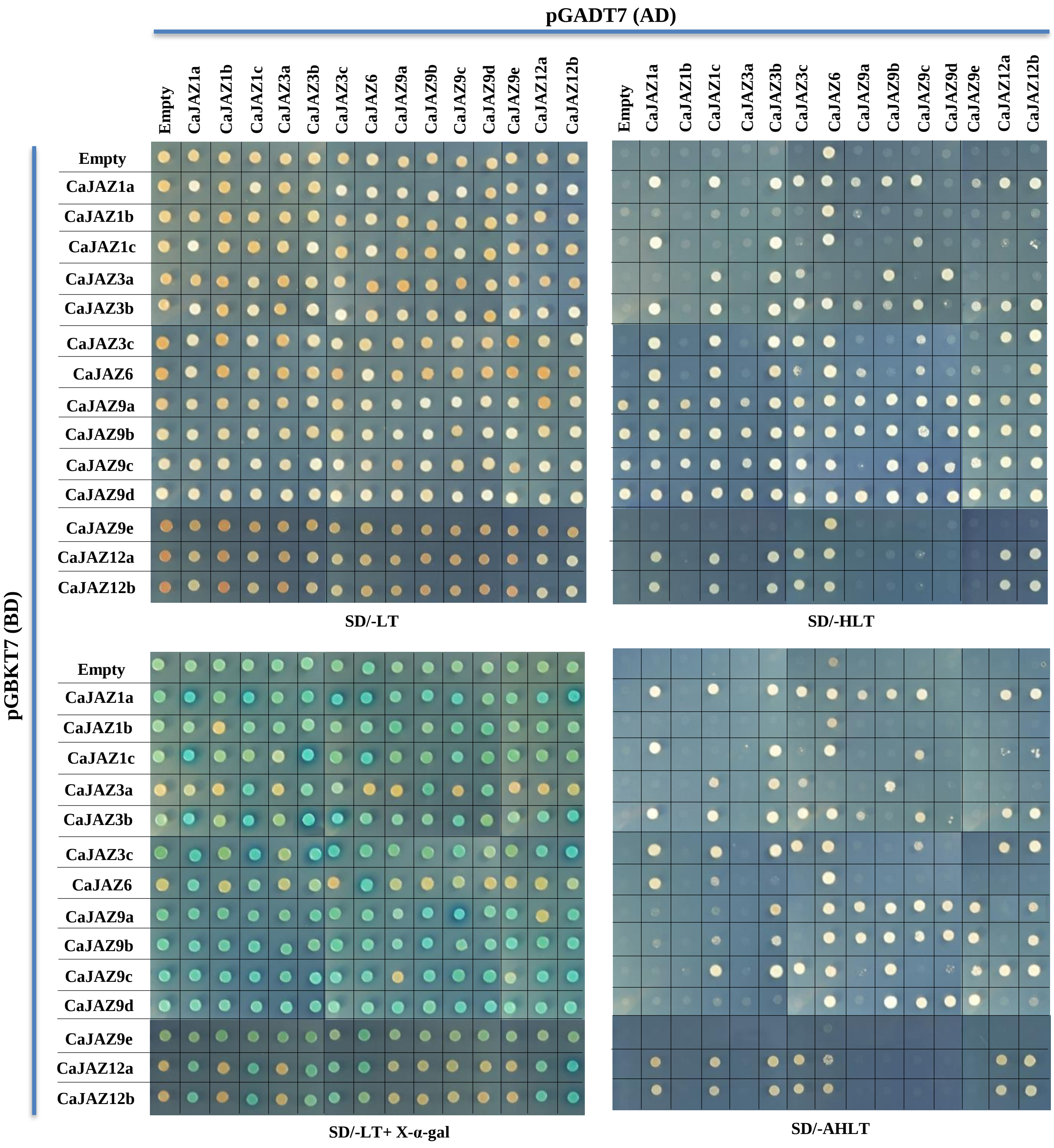


**Figure S9. CaJAZ proteins interacts with each other to form homo and hetero-dimers.** Double transformants having respective pGADT7 and pGBKT7 clones were inoculated on SD-LT, SD-HLT, SD-LT supplemented with X-alpha-Gal (20 mg/L) and SD-AHLT dropout media. Observations were recorded after 3 days of incubation at 30°C.

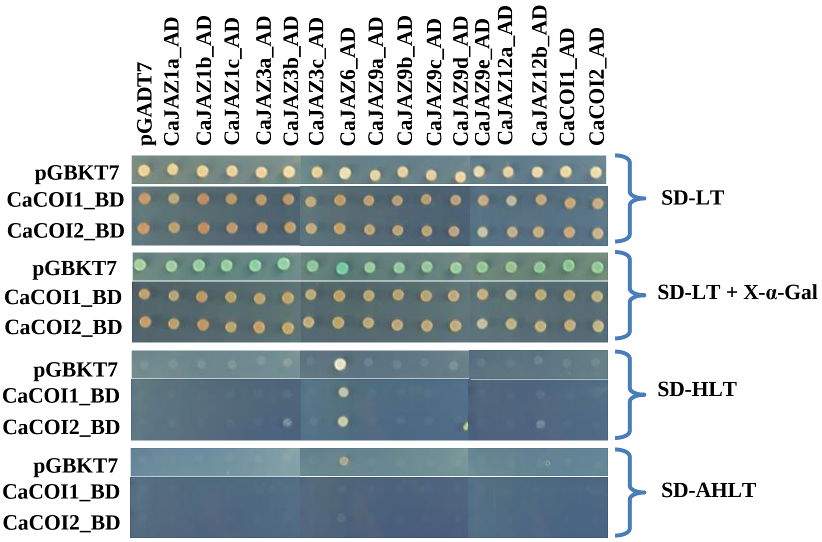


**Figure S10. CaCOI proteins fail to interact with CaJAZ proteins in the absence of coronatine.** Double transformants having respective pGADT7 and pGBKT7 clones were inoculated on SD-LT, SD-LT supplemented with X-alpha-Gal (20 mg/L), SD-HLT and SD-AHLT dropout media. Observations were recorded after 3 days of incubation at 30°C.

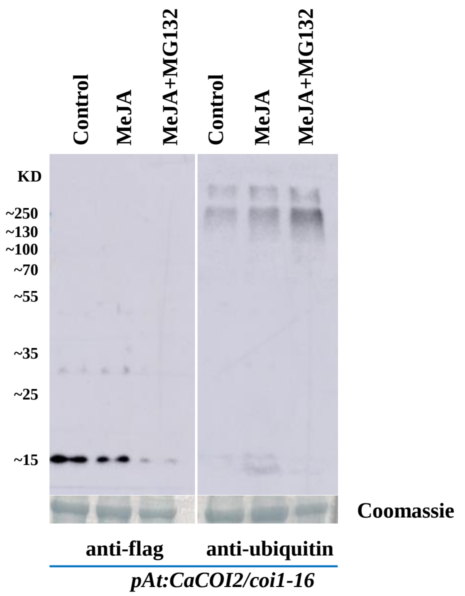


**Figure S11. In-vivo ubiquitination pattern in *pAt:CaCOI2* complemented *coi1-16* lines.** Total protein was isolated from pAt:CaCOI2 complemented lines during control, 100µM MeJA or 100µM MeJA and 50µM MG132, respectively. ~10µg protein was loaded to each well. Coomassie staining indicates equal loading.

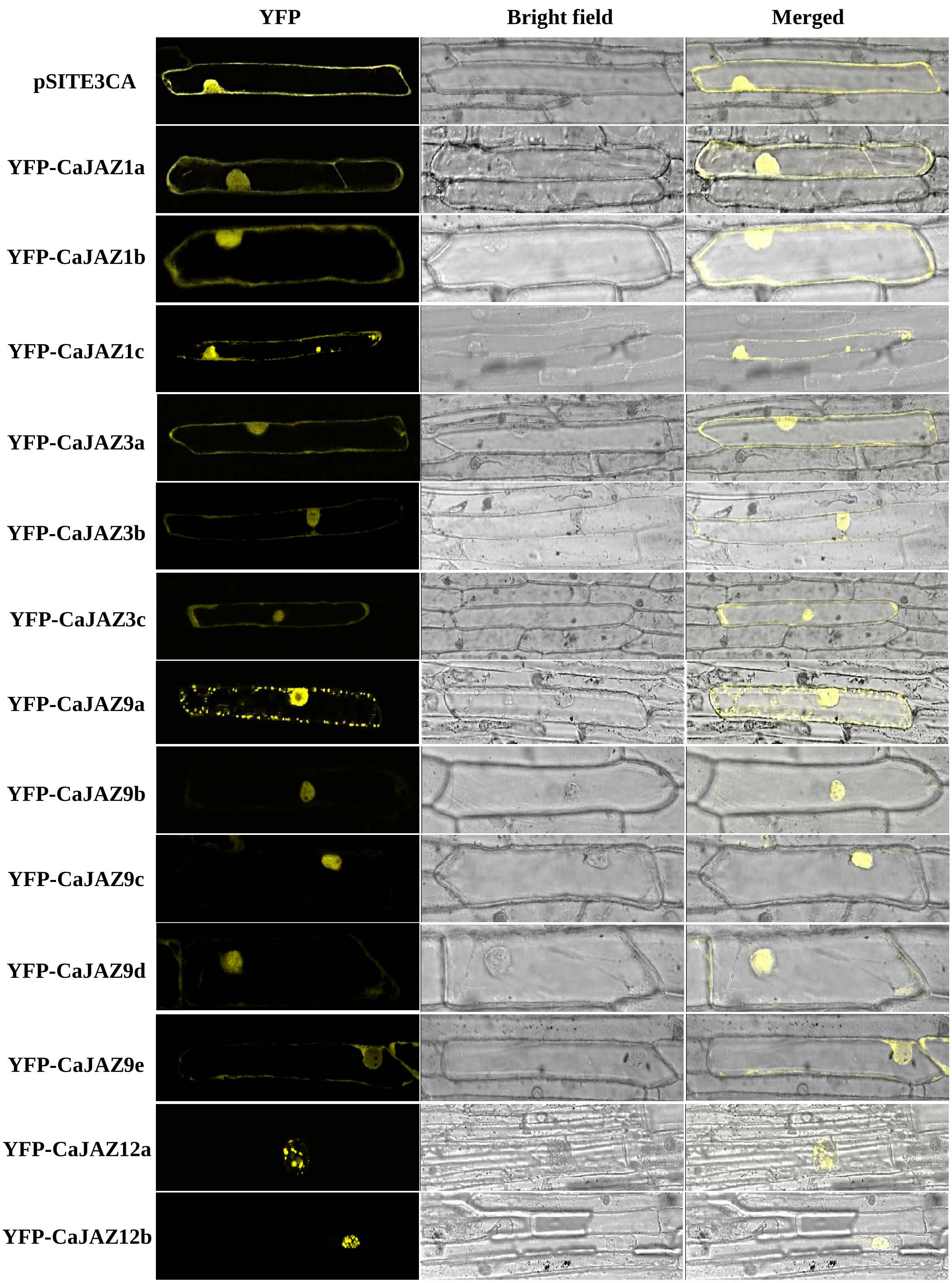


**Figure S12. Subcellular localization of identified JAZ proteins.** Longest ORF was amplified and cloned in pSITE3CA vector to obtained YFP-gene fusion constructs. The respective constructs were transformed in onion epidermal cells using gold particle bombardment and visualised the YFP signals under confocal microscope (Leica SP8) second day after bombardment .

```
AtCOI1    M-EDPDIKRCKLSCVATVDDVIEQVMTYITDPKDRDSASLVCRRWFKIDSETREHVTMAL    59
CaCOI1    MEEKAACGGVGRTSTKLSDVVLDCVMPYIHDPKDRDAVSQVCRRWYELDSLTRKHVTIAL    60

          [coi1-5, G98D]
AtCOI1    CYTATPDRLSRRFPNLRSLKLKGKPRAAMFNLIPENWGGYVTPWVTEISNNLRQLKSVHF    119
CaCOI1    CYTTTPDRLRRRFPHLESLKLKGKPRAAMFNLIPENWGGFVTPWVKEISKYFDCLKSLHF    120

          [coi1-7, G155E]
AtCOI1    RRMIVSDLDLDRLAKARADDLETLKLDKCSGFTTDGLLSIVTHCRKIKTLLMEESSFSEK    179
CaCOI1    RRMIVTDSDLQILARSRCNSLHALKLEKCSGFSTDGLYYVGRFCKNLRVLFMEESSVVEK    180

AtCOI1    DGKWLHELAQHNTSLEVLNFYMTEFAKISPKDLETIARNCRSLVSVKVGDFEILELVGFF    239
CaCOI1    DGEWLHVLALNNTVLETLNFYLTDIANVRIQDPELIAKNCPNLVSVKITDCEILNLMNFF    240

          [coi1-16, L245F; coi1-2, L245F]
AtCOI1    KAAANLEEFCGGSLNEDIGMPEKYMNLVFPRKLCRLGLSYMGPNEMPILFPFAAQIRKLD    299
CaCOI1    RYASSLEEFCGGSYNED---PEKYSAISLPAKLSRLGLTYIGKNEMPFVFPYAAMLKKLD    297

          [coi1-21rsp, G330E] [coi1-6, Q343*]
AtCOI1    LLYALLETEDHCTLIQKCPNLEVLETRNVIGDRGLEVLAQYCKQLKRLRIERGADEQGME    359
CaCOI1    LLYAMLDTEDHCTLIQKCPNLEVLESRNVIGDRGLEVLASCCKKLRRLRIERGDDDQGME    357

          [coi1-4, G369E] [coi1-32, G399D]
AtCOI1    DEEGLVSQRGLIALAQGCQELEYMAVYVSDITNESLESIGTYLKNLCDFRLVLLDREERI    419
CaCOI1    DEEGIVSQRGLIALSQGCPELEYMAVYVSDITNASLEHIGTHLKNLCDFRLVLLDREEKI    417

          [coi1-22rsp, G434E] [coi1-34, A442T] [coi1-9, D452A] [coi1-1, W467*]
AtCOI1    TDLPLDNGVRSLLIGCKKLRRFAFYLRQGGLTDLGLSYIGQYSPNVRWMLLGYVGESDEG    479
CaCOI1    SDLPLDNGVRALLRGCDKLRRFALYLRPGGITDVGLGYIGQYSPNVRWMLLGYVGETDAG    477

          [coi1-10, L490A]
AtCOI1    LMEFSRGCPNLQKLEMRGCCF-SERAIAAAVTKLPSLRYLWVQGYRASMTGQDLMQMARP    538
CaCOI1    LLEFSKGCPSLQKLEMRGCSFFSEYALAIAATRLTSLRYLWVQGYGASPSGRDLLAMARP    537

          [coi1-8, E543K]
AtCOI1    YWNIELIPSRRVPEVNQ-QGEIREMEHPAHILAYYSLAGQRTDCPTTVRVLKEPI-----    592
CaCOI1    YWNIELIPSRRVVVKNQ-QDELVAVEHPAHILAYYSLAGPRSDFPDTVIPLDPAAYY---    593

AtCOI1    ---   592
CaCOI1    ---   593
```

**Figure S13. Multiple alleles are associated with the functionality of COI1 as JA receptor** (Feys et al., 1994; Ellis and Turner, 2002; Yan et al., 2009; Acosta et al., 2013; Song et al., 2021).

```
AtCOI1          MEDPD-I---------KRCKLSCVATVDDVIEQVMTYITDPKDRDSASLVCRRWFKIDSE   50
CaCOI1          MEEKAAC---------GGVGRTSTKLSDVVLDCVMPYIHDPKDRDAVSQVCRRWYELDSL   51
CaCOI2          MT--------------EDRSVKTNRVVDVVLDCVIPYIDDPKDRDAVSQVCRRWYELDSL   46
ZmCOI1a         MGGEVP-EPRRLSRALSFG---CGGVPEEALHLVFGYMDDPRDREAASLVCRLWHRIDAL   56
ZmCOI1b         MGGEAP-EPRRLTRALSIGGGDGGWVPEEMLQLVMGFVEDPRDREAASLVCHRWHRVDAL   59
ZmCOI1c         MGGEAP-EPRRLTRALSIGGGGGGWVPDEMLQLVLGFVEDPRDREALSLVCRRWHRIDAL   59
ZmCOI1d         MGGEVP-EPRRLSRSLSFG---CGGVPEEALHLVLGYVDDPRDREAASLVCRRWHRIDAL   56
ZmCOI2a         MGGEAEGGERRLGRVLSFG------IPDTALGLVMGYVEDPWDRDAISLVCRHWCRVDAL   54
ZmCOI2b         MGGEAEGGERRLGRVLSFG------IPDTALGLVMGYVEDPWDRDAISLVCRHWCRVDAL   54
OsCOI1a         MGGEVP-EPRRLNRALSFD----DWVPDEALHLVMGHVEDPRDREAASRVCRRWHRIDAL   55
OsCOI1b         MGGEAP-EARRLDRAMSFGG--AGSIPEEALHLVLGYVDDPRDREAVSLVCRRWHRIDAL   57
OsCOI2          MGG--EAGERRLGRAMSFG------IPDVALGLVMGFVEDPWDRDAISLVCRHWCRVDAL   52
                                           :  :  *: .: ** **:: * **: * .:*:
```

**Figure S14. Amino acid sequence alignment showing conserved acidic amino acid at (AtCOI1$^{T29}$) position.**

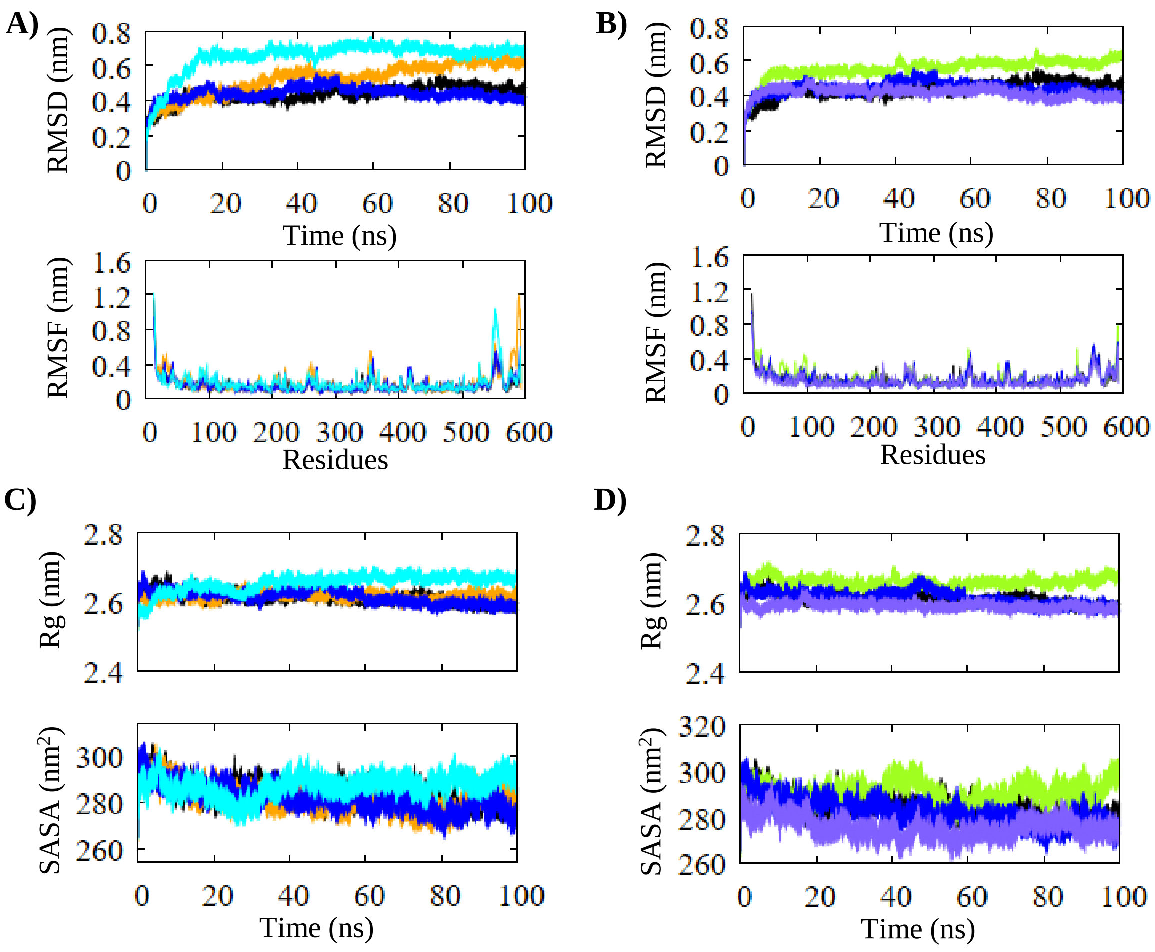


**Figure S15. Docking analysis of AtCOI1 with InsP8 and JA-Ile.** RMSD and RMSF graphs for AtCOI1-InsP8 **(A)** and AtCOI1-JA-Ile **(B)**. Rg and SASA graphs for AtCOI1/InsP8 **(C)** and AtCOI1/JA-Ile **(D)**. Colour scheme for graphs is as follows: AtCOI1$^{T29}$ (Black), AtCOI1$^{H29}$ (Blue), AtCOI1$^{T29}$/InsP8 complex (Orange), AtCOI1 $^{H29}$/InsP8 complex (Cyan), AtCOI1 $^{T29}$/JA-Ile (Lime), AtCOI1 $^{H29}$/JA-Ile (Slate).
